# Lower air humidity reduced both the plant growth and activities of photosystems I and II under prolonged heat stress

**DOI:** 10.1101/2022.06.03.494694

**Authors:** Eugene A. Lysenko, Marina. A. Kozuleva, Alexander A. Klaus, Natallia L. Pshybytko, Victor V. Kusnetsov

## Abstract

The warming is global problem. In natural environments, a heat stress is accompanied with a drought usually. The effect of lower air humidity remains obscure. Maize and barley plants were supplied with an unlimited source of water for the root uptake and undergone to heat stress for 48 h at contrast conditions of air humidity. The lower air humidity decreased photochemical activities of photosystem I and photosystem II. The small effect was revealed in control. The temperature elevation to 37°C and 42°C increased relative activities of the both photosystems; the photosystem II was activated more. The effect of air humidity disappeared at 37°C; at 42°C, the effect was small. At 46°C, lower air humidity magnified substantially the inhibitory effect of heat. Consequently, the maximal and relative activities of the both photosystems were decreased in maize and barley; the plant growth was reduced greatly. The photosystem II was inhibited more. At 46°C, maize plants increased water uptake by roots at lower air humidity and survived; barley plants were unable to increase water uptake and died. Therefore, air humidity is the important component of environmental heat stress influencing activities of photosystem I and photosystem II and thereby plant growth and viability.

**Highlight:** The effect of severe heat stress was magnified with lower air humidity. At 46°C, their mutual action inhibited photosystem I and II and reduced greatly the plant growth and viability.

## Introduction

High temperature is one of the major factors influencing the growth and development of plants. The temperature increase by 10-15°C above habitual is considered as heat stress or heat shock (Wahid *et al*., 2007). The process of global warming increases the rate of heat stress (HS) conditions in the natural environment. The productivity of major agronomic cultures is in a bona fide correlation with the environmental changes of temperature; the increase of temperature in growing season is accompanied with about 17% decrease of corn and soybean yield for the each Celsius degree increase (Lobell and Asner, 2003). The photosynthesis is considered as one of the most heat sensitive function in plant organisms (Berry and Björkman, 1980). The reduction of photosynthesis can underlay the decrease of plant growth and yield (Sharkey, 2005).

The impact of elevated temperature on photosynthesis was studied for decades. The most important targets of HS are Rubisco activase in Calvin-Benson cycle, oxygen evolving complex and D1 protein in the photosystem II (PSII), and ATP synthesis (reviewed in Sharkey, 2005; Allakhverdiev *et al*., 2008). The elevation of temperature increases the fluidity of thylakoid membranes (Horvath *et al*., 1998); in turn, this leads to the unstacking and disorganization of the membranes (Gounaris *et al*., 1984). High temperature or membrane state induces migration of PSII antenna complexes to photosystem I (PSI) (Pastenes and Horton, 1996). The state transition induced with heat acts through same mechanism as high light: LHCII phosphorylation with the kinase STN7 is the key step in this process; however, heat induced state transition can be found in the darkness (Nellaepalli *et al*., 2011). The lack of trienoic fatty acids in the thylakoid membranes decreases their fluidity and increased plant tolerance to heat (Murakami *et al*., 2000); saturation of membrane lipids saved activity of PSII at elevated temperature in green alga (Sato *et al*., 1996).

An elevated temperature is combined with two other factors usually. Naturally, a rise of temperature is caused with the sun light; therefore, high light and high temperature conditions are overlapped frequently. Many investigations were devoted to simultaneous application of heat and high light stresses to the photosynthetic apparatus of plants (*e.g*., Monneveux *et al*., 2003; Diaz *et al*., 2007; Zhou *et al*., 2020). In turn, elevated temperature increases water evaporation; therefore, heat and drought stresses coincide in many habitats. Last years, cooperative action of heat and drought has been realized. For example, impacts of HS and drought on maize plants were reviewed recently (Chavez-Arias *et al*., 2021). However, studies of heat and drought impact on photosynthetic processes remains to be scarce. It was shown that heat and drought decreased the stomatal conductance, transpiration, and gas exchange with the concomitant decrease of CO_2_ assimilation rate (Killi *et al*., 2017, 2020; Alhaithloul, 2019). Activity of PSII was impaired also: both the potential (Fv/Fm) and actual (Φ_PSII_) quantum yields were decreased in the conditions of HS and drought; maximal quantum yield in dark adapted state (Fv/Fm) was inhibited to a lesser extent compared with large decrease of quantum yield in light adapted state (Φ_PSII_) (Killi *et al*., 2017, 2020; Alhaithloul, 2019; Yudina *et al*., 2020). All the abovementioned articles demonstrated the increase of non-photochemical quenching of light energy in the conditions of HS and drought. In pea plants, both the HS and drought stimulated the increase of non-photochemical quenching; however, at the same conditions HS did not change non-photochemical quenching in pumpkin and decreased significantly non-photochemical quenching in wheat (drought impact was not studied for pumpkin and wheat) (Yudina *et al*., 2020).

In all these studies, the effect of drought was caused by the restriction of water uptake by roots. This experimental design demonstrates effect of water lack in soil. However, the specific feature of arid climate is low water content in both soil and air. The temperature increases were accompanied with the decreases of air relative humidity (RH) (Arena *et al*., 2020). Lower RH of air should stimulate gas exchange between leaves and environment. Such an increase gives plants two advantages and one disadvantage. First, increased gas exchange causes higher CO_2_ uptake by leaves that is beneficial for the photosynthesis and plant growth. Second, increased gas exchange resulted in more water evaporation from leaves. Water evaporation enables to decrease temperature of a leaf blade. Under an elevated temperature, plants are able to decrease leaf temperature by several degrees of Celcius grade comparing with air temperature (Wise *et al*., 2004; Camejo *et al*., 2005). Transpirational cooling of leaves is considered as important mechanism of plant adaptation to HS (Radin *et al*., 1994; Sharkey, 2005). The disadvantage appears the reverse side of water evaporation: leaves losses more water.

Air humidity may influence processes including such unexpected as wound sealing (Speck *et al*., 2018). The role of air humidity in the action of HS remains unclear; its effect on the photosynthesis is unexplored. The understanding of air humidity impact is important for the both environmental and laboratory studies exploring the HS influence on the photosynthesis. Most studies of photosynthesis under HS omitted the information concerning air humidity. We found the only work explained that control plants were kept at 70±7% RH and experimental plants were heated at 48±6% RH (Sharma *et al*., 2015). Before the HS experiments, plants were grown in the conditions varied from 40% RH (Pastenes and Horton, 1996; Doğru, 2021) to ~90% RH (Arena *et al*., 2020); more often plants were grown at the conditions 60-65% RH (Havaux, 1996; Yan *et al*., 2013; Turan *et al*., 2019; Ferguson *et al*., 2020; Yudina et al *et al*., 2020). We are unsure is it critical or not. Is it correct to compare data obtained at diverse RH background? It will be beneficial to carry out an experiment with plants having unlimited source of water for the root uptake and undergone to HS at contrast levels of air humidity.

The experimental design varies much in the numerous studies of HS action on the photosynthesis. Two main factors are the temperature level and time of elevated temperature action. The temperature range 35-40°C is considered as moderate heat stress (Sharkey, 2005); the heat shock requires the temperature 40-42°C and higher. The temperature above 45°C induces irreversible changes in PSII activity and can be lethal (Sharkey, 2005). The lethal effect can be avoided with short time of temperature elevation.

The time of the HS treatment varies much. Short time application is very popular. In many studies HS was applied for 5 min (Feller *et al*., 1998), 10 min (Turan *et al*., 2019), 15 min (Fergusson *et al*., 2020), 20 min (Doğru, 2021), and 30 min (Yudina *et al*., 2020). Short time HS (5-15 min) was obtained by the immersion of leaves into water with the corresponding temperature; HS treatment in hot air started from 20 min. The range of water-induced HS varied from 1 min (Havaux and Strasser, 1990) to 80 min (Havaux *et al*., 1991). The brief application is focused on a direct harm or alteration of the photosynthetic molecular complexes; this approach is the most popular among photobiologists. The longer application of HS involves in the analyses protective mechanisms of cellular level, for example, synthesis *de novo* of photosynthetic polypeptides that requires all the steps of gene expression from transcription (Zubo *et al*., 2008) to translation (Franco *et al*., 1999; Tanaka *et al*., 2000). The prolongation of HS until few hours - 1h (Arena *et al*., 2020), 2h (Camejo *et al*., 2005; Yan *et al*., 2013), and 3h (Kalituho *et al*., 2003) – is less popular. The application of HS for days or many hours at least allows studying protective mechanisms at the level of a whole plant. The long experiments are rare. We has found single investigations studied HS effect on the photosynthesis for 6 h (Wang *et al*., 2020), one week (Scharma *et al*., 2015), and two weeks and more (Killi *et al*., 2020).

The studies mentioned above used for the HS analysis single temperature or wide range with the gradual increase of temperatures. Some articles demonstrated plain and sufficient experimental design. The photosynthesis was studied in the conditions of moderate HS (35/38°C), heat shock or genuine HS (40/43°C) and damaging HS (45/48°C) (Feller *et al*., 1998; Yan *et al*., 2013). However, HS experiment is restricted by the simple rule: The higher temperature the shorter time. The highest temperatures 50-55°C were applied for 5-15 min only (Havaux, 1996; Bukhov and Carpentier, 2000; Turan *et al*., 2019; Fergusson *et al*., 2020). The temperature 45°C was applied for a couple of hours (Camejo *et al*., 2005; Yan *et al*., 2013); the treatments for days or 6h were restricted to 35-38°C (Scharma *et al*., 2015; Killi *et al*., 2020; Wang *et al*., 2020). It is known that light increased the resistance of photosynthesis to HS during the short or moderate time of treatment (Havaux and Strasser, 1990; Havaux *et al*., 1991; Kalituho *et al*., 2003). Probably, continuous illumination will enable applying temperatures above 40°C for few days.

The current work has analyzed the influence of air humidity on the plant growth and activities of PSI and PSII under an elevated temperature. The analyses comprised heat tolerant (*Zea mays*) and heat sensitive (*Hordeum vulgare*) plant species belonging to Poaceae family. Hydroponically grown seedlings were placed in thermostat chambers for 48 h; the seedlings were supplied with the unlimited source of water for root uptake. The both plant species were grown 48 h under the conditions of control (24°C), moderate HS (37°C), genuine HS (42°C), and nearly lethal HS (46°C). The each temperature treatment was performed in parallel at higher and lower relative humidity of air. The effects of HS and air RH on plant growth and photochemical activities of PSI and PSII were studied. The measurement of chlorophyll (Chl) fluorescence and P_700_ light absorption was carried out at the temperature of treatment (24-48°C, correspondingly). The data obtained are presented below.

## Methods

### Plant growth conditions

Maize (*Zea mays* L. cv. Luchistaya) and barley (*Hordeum vulgare* L. cv. Luch) seedlings were grown in phytotron chambers at 180-200 μmol photons m^−2^ s^−1^ and a photoperiod of 16 h light/8 h dark under continuous aeration on modified Hoagland medium (Lysenko *et al*., 2019). The temperature was optimal for each species: 25-26°C for maize and 21°C for barley. Caryopses were immersed in 0.25 mM CaCl_2_ for 30 min and then kept for 2 days at 4°C in the dark on a filter paper moistened with 0.25 mM CaCl_2_. The caryopses were placed in growth conditions, and the plant age was determined from this time. Two days later, the seedlings were transferred to pots with Hoagland medium.

### Plant growth at control or elevated temperatures (experiment)

Pots with seven-day-old seedlings were transferred to thermostat chambers for 48 h; they were grown under continuous illumination at 60-80 μmol photons m^−2^ s^−1^. Each of the four temperature regimes was accomplished in parallel at two different levels of relative air humidity (Table 1); it was achieved in thermostats of two types. Thermostats of both types used similar heating elements and different cooling elements. The higher humidity (HH) conditions were kept in a thermostat with a ventilation system (TS-1/80 SPU, “Smolenskoe SKTB SPU”, Russia). The lower humidity (LH) conditions were achieved in a thermostat with a refrigeration unit (TSO-200 SPU, “Smolenskoe SKTB SPU”, Russia); water vapor was condensed on the chilled elements of the refrigeration unit therefore air humidity was decreased in the chamber largely. The both cooling systems were outside of the chambers with plants. The conditions of air humidity were measured with the use of a hygro-thermometer AZ 8721 (AZ Instrument, China) (see Table 1).

**Table.**
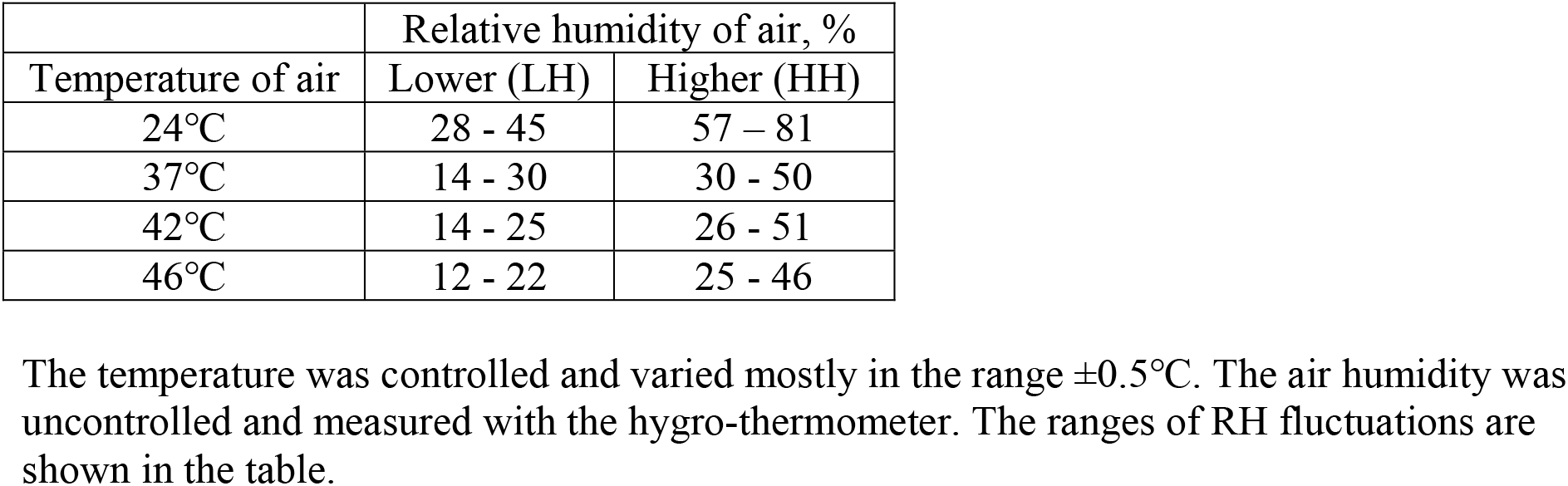
The temperature and relative humidity of air in the thermostat chambers during experiment.

| Temperature of air | Relative humidity of air, % |  |
| --- | --- | --- |
|  | Lower (LH) | Higher (HH) |
| 24°C | 28 - 45 | 57 – 81 |
| 37°C | 14 - 30 | 30 - 50 |
| 42°C | 14 - 25 | 26 - 51 |
| 46°C | 12 - 22 | 25 - 46 |

The temperature was controlled and varied mostly in the range $\pm 0.5^\circ\text{C}$ . The air humidity was uncontrolled and measured with the hygro-thermometer. The ranges of RH fluctuations are shown in the table.

In the beginning of temperature treatment, each pot with plants was filled with Hoagland medium till the pot’s edge. New portions of the media were added several times per a day including at the end of experiment. All the portions added after the beginning were summarized and divided per number of plants in a corresponding pot; each pot bore 25±1 plants. The average water uptake by one plant during 48 h was calculated and used to estimate the total water evaporation by plants.

Plants of the both species were grown simultaneously and exposed to a one of the four temperatures under the both levels of air RH. The analysis with the method of pulse amplitude modulation (PAM) lasted whole day. The exposition to particular temperature was initiated in a morning for one species and afternoon for another species. PAM analysis was started in a morning or afternoon, correspondingly. Thus, plants were undergone to each particular temperature for 48±3 h before the measurement of PSI and PSII activities. In the all except one variant of treatment, plants were alive at the end of the experiment. Under conditions 46°C at lower air RH, barley plants were dying fast after 51-52 h of treatment.

For each variant of *in vivo* treatment (48 h), biological experiments were repeated four times.

### Temperature background of PAM analysis

The maize and barley plants were adapted to the darkness for 30 min in the thermostat (TS-1/80 SPU). The dark-adaptation was carried out at the same temperature that was applied to plants in the previous 48 h (24°C, 37°C, 42°C, or 46°C correspondingly). The dark-adapted plants were transferred rapidly to the Dual-PAM-100 (Walz, Germany), avoiding bright light, and kept in the dark for another 4 min. The leaves were inserted between two measuring heads of the Dual-PAM-100; the measuring heads mounted to stand were kept in the thermostat chamber also (TS-1/80 SPU). The extra adaptation to darkness (4 min) and all the processes of measurement were performed at the temperature corresponding to the temperature of treatment in previous 48 h (24°C, 37°C, 42°C, or 46°C correspondingly). The roots of plants measured were submerged in Hoagland medium that was critical at higher temperatures.

Plants were preadapted to darkness (30 min) and measured (~20 min) at the temperature of whole treatment (48 h) to save all possible adaptations developed with plants during two days at HS. The chambers were opened too frequently to save specific air humidity during adaptation to darkness and measurement; unavoidably, it should be ambient. Therefore single type of thermostat was used for the plants adapted to the both HH and LH conditions.

### PAM analysis

The chlorophyll a fluorescence and P_700_ light absorption were registered simultaneously with the Dual-PAM-100. The largest fully developed leaf was used for the measurement: the 2^nd^ leaf of maize and the first leaf of barley. The Chl a fluorescence was excited at 460 nm (9 μmol photons m^−2^ s^−1^; the measuring light). P_700_ is the reaction center chlorophyll of PSI; the level of oxidized P_700_ was measured as the difference in light absorption at 830 and 875 nm. The red light 635 nm was used as the actinic light (AL) and for the saturation pulses (SPs).

In the dark, the measuring light only was used for the determination of minimal (Fo) Chl fluorescence; the first SP was applied for the measurement of maximal (Fm) Chl fluorescence; the pre-illumination with far-red light (720 nm, Int. #10) followed with the second SP were employed for the determination of minimal (Po) and maximal (Pm) P_700_ absorption.

Next, the AL was turned on and adaptation of electron-transport chain to the light condition was studied; this method is called “induction curve” (IC). The intensity of AL (70 μmol photons m^−2^ s^−1^) corresponded to the illumination level in the thermostat chambers during experiment.

The AL was applied for 7.5 min; SPs (4 mmol photons m^−2^ s^−1^, 500 ms) were induced every 40 s and followed by far-red light (Int. #10) for 5 s. The maximum level of fluorescence in the light (Fm′) and maximal P_700_ change in the light (Pm′) were measured with SPs. The minimum level of fluorescence in the light (Fo′) was determined with the far-red illumination. The steady state Chl fluorescence (Fs) and P_700_ absorption were measured at the end of each 40s period.

The “induction curve” method is the correct and, probably, preferable way of plant adaptation to light conditions before an application of “rapid light curve” (RLC) method (Lysenko, 2021). After the end of IC measurement, the measurement of the RLC was started immediately. In RLC, each step of actinic light lasted for 30 s; the set of AL intensities is shown in the figures. The other parameters were the same as in IC.

The quantum yield of PSI was calculated as follows: Y(I) = (Pm′ – P)/(Pm–Po) (Klughammer and Schreiber, 1994). The parameters of Chl fluorescence were calculated using the following equations: Fv = Fm – Fo, Fv′ = Fm′ – Fo′, qP = (Fm′ – Fs)/Fv′ (Schreiber *et al*., 1986; van Kooten and Snel, 1990); Φ_PSII_ = (Fm′ – Fs)/Fm′ (Genty *et al*., 1989; Kalaji *et al*., 2014); X(II) = (Fm′ – Fs)/Fv (Lysenko *et al*., 2020).

### Chlorophyll estimation

In a leaf blade, the terminal 1-cm segment was removed, and the following 1-cm segment was used for the analysis. Chlorophylls and carotenoids were extracted with 80% acetone, concentrations were estimated in mg units according to (Lichtenthaler, 1987).

### Statistics

The data were processed using the Excel (Microsoft) software. The significance of differences between mean values was verified using the two-tailed Student’s t-test.

In few particular cases, the significance of differences between steady state levels of IC was verified additionally using the z-test (Supplementary Commentary 1 at *JXB* online) or paired nonparametric binomial test (Supplementary Commentary 2).

## Results

### Plant growth

The set of elevated temperatures imposed diverse effects on the plant growth (Fig 1, Supplementary Tables S1-S5). The moderate HS (37°C) had no obvious effect on the growth of maize shoots; maize roots accumulated less weight (significant at LH conditions). Barley second leaves were growing fast; their growth was inhibited at 37°C according to all the parameters studied (Fig 1H,J; Supplementary Table S3). Because of this, the height and FW of barley shoots were diminished (Fig 1D; Supplementary Table S2). The genuine HS (42°C) inhibited growth of the both barley and maize seedlings; the inhibitory effect was significant for many parameters studied (Fig 1, Supplementary Tables S1-S5). The nearly lethal HS (46°C) retarded the growth more. The drastic decreases of size, weight, and water content (see below) were detected at 46°C only. In such cases, the LH conditions reduced size and weight to a larger extent; it was observed in the both species. Rarely, the HH and LH conditions imposed an equal effect (e.g., maize root FW) (Fig 1, Supplementary Tables S1-S5). Under the conditions 46°C at LH, barley plants died immediately when the experimental treatment exceeded 48 h for three-five hours; therefore, many parameters were not determined in this variant.

**Fig. 1.**
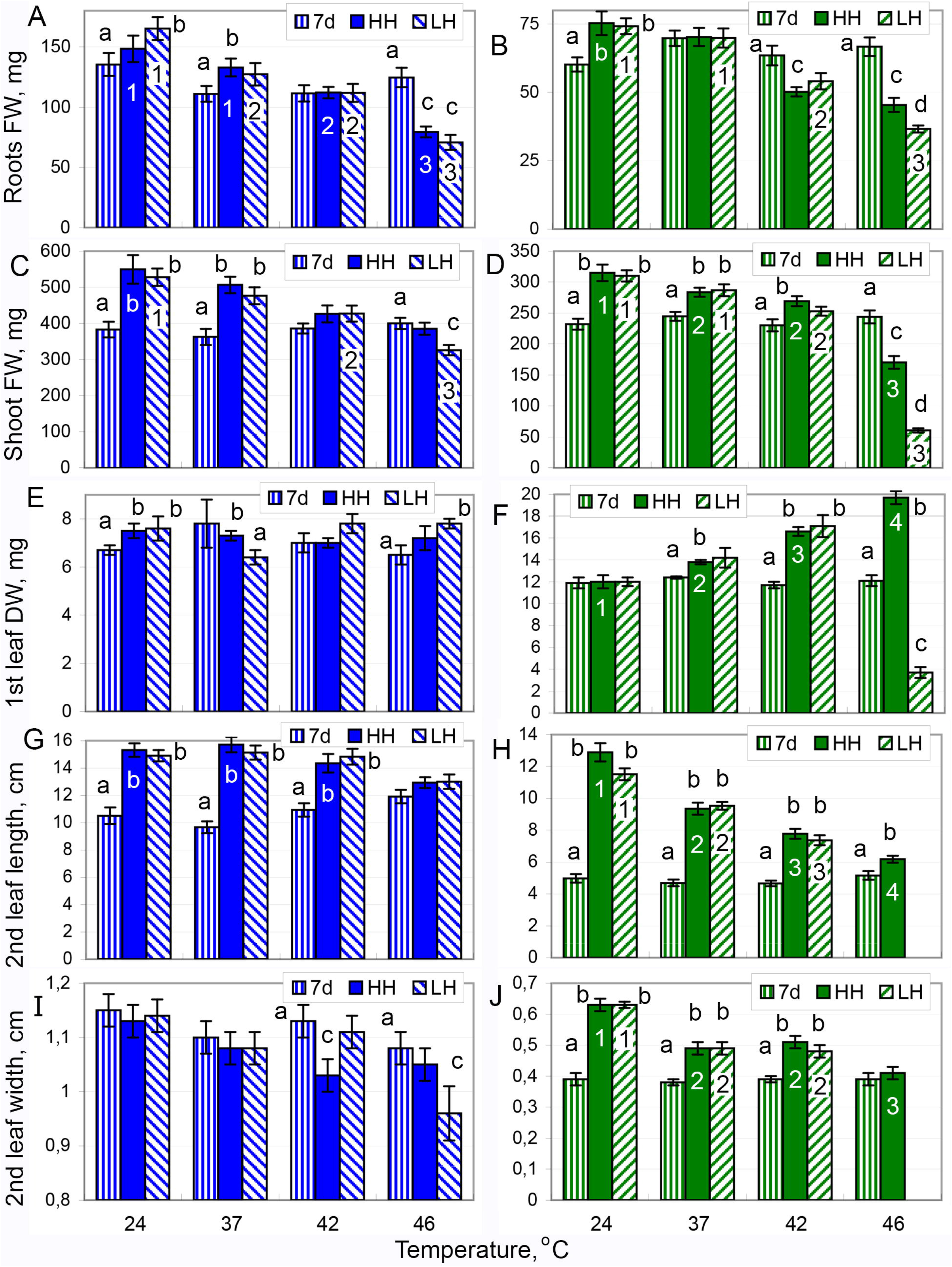
Effect of elevated temperature in contrast RH conditions on the growth of maize and barley plants. The weight and size of plants before the experiment (7d, vertically hatched bars) and after 48 h exposure to HS at the contrast RH conditions (HH, filled bars, or LH, diagonally hatched bars). A, B – root FW; C, D – shoot FW; E, F – first leaf DW; G, H – second leaf length; I, J – second leaf width. A, C, E, G, I – maize; B, D, F, H, J – barley. Data are presented as means ± standard error (SE). 1-4 - differences between variants of temperature regime are significant at p ≤ 0.05. a-d - differences between plants at same temperature regime before (7d) and after experiment at different RH conditions (HH & LH) are significant at p ≤ 0.05. The results of statistical analysis are shown partially to ease understanding of the figure. For the effect of RH (a-d), statistically different values are marked only. For the effect of temperature (1-4), the most interesting examples of gradual decline or rise are shown only. The full dataset of weight and size parameters and complete statistical analysis are given in Supplementary Tables S1-S5.

For many parameters, a gradual retardation effect was found; the stepwise elevation of temperature caused consecutive reduction of size or weight (Fig 1, Supplementary Tables S1-S5).

In 7-day old seedlings, the first leaves completed their growth mostly and second leaves were growing actively. Two days later, the second leaves were fully developed in maize and still growing in barley. The elevation of temperature for 48 h slowed down the growth or even reduced size or weight comparing with the initial 7-day old plants. Few cases of reduction were found at the temperature 42°C. The barley roots lost FW and the maize leaves reduced their widths (Fig 1B,I, Supplementary Tables S1, S3, S4). At 42°C, the losses of weight or size were relatively small. Under the temperature 46°C, plants lost FW of the both roots and shoots. It was revealed in the both species at the both conditions of air humidity (Fig 1). The loss of FW by roots was mainly due to the loss of DW (Supplementary Table S1); the loss of FW by shoots was caused mostly by water loss (see below). Some organs of shoots lost DW significantly (Fig 1F); however, there were loss of DW by neither whole shoots nor whole seedlings. At 46°C, the losses of weight were large especially in barley and at LH conditions. Under the temperature 46°C at LH, barley plants were reduced greatly comparing with the control: DW of the first leaves was reduced three times, FW of shoots was decreased four times, and FW of the first leaves was reduced five times (Supplementary Tables S1-S5). Under the temperature 46°C, plant leaves became to be narrowed; the first leaves of barley and maize and second leaves of maize were reduced in widths by 1 mm (Fig 1I,J, Supplementary Table S3-S4).

We revealed a paradoxical reaction: the elevation of temperature stimulated the gradual increase of DW in the first leaves of barley (Fig 1F). At this developmental stage, the first leaf is main organ of barley seedling. Probably, barley seedlings focused transport of organic and/or mineral substances to the first leaves for surviving in stressful conditions. In the most severe environment (46°C at LH), barley seedlings were unable to overcome unfavorable conditions, lost DW, and died few hours later.

### Water content and uptake

The water content in the both species decreased at 37°C; the further elevation of temperature induced gradual decrease of water content in tissues (Fig 2. Supplementary Table S6). The roots lost small portion of water even at the highest temperature; probably, because roots were immersed in water. The stem (with leaf sheaths) demonstrated small loss of water also (Supplementary Table S6). In the leaves, water content was reduced substantially or even dramatically; this resulted in the large water loss by the whole shoots. Barley plants lost more water comparing with maize plants. The air humidity did not influence water content at the temperatures 24-42°C with a single exception. In barley plants grown at 42°C, the water loss in stem was larger in LH environment than in HH environment; however, the difference was small (Supplementary Table S6). Under the nearly lethal HS (46°C), the LH conditions stimulated larger water loss comparing with HH conditions (Fig 2. Supplementary Table S6). The opposite tendency was revealed only in the first leaves of barley (Fig 2F). In this case, alterations of water were masked with the changes of DW (see Discussion).

**Fig. 2.**
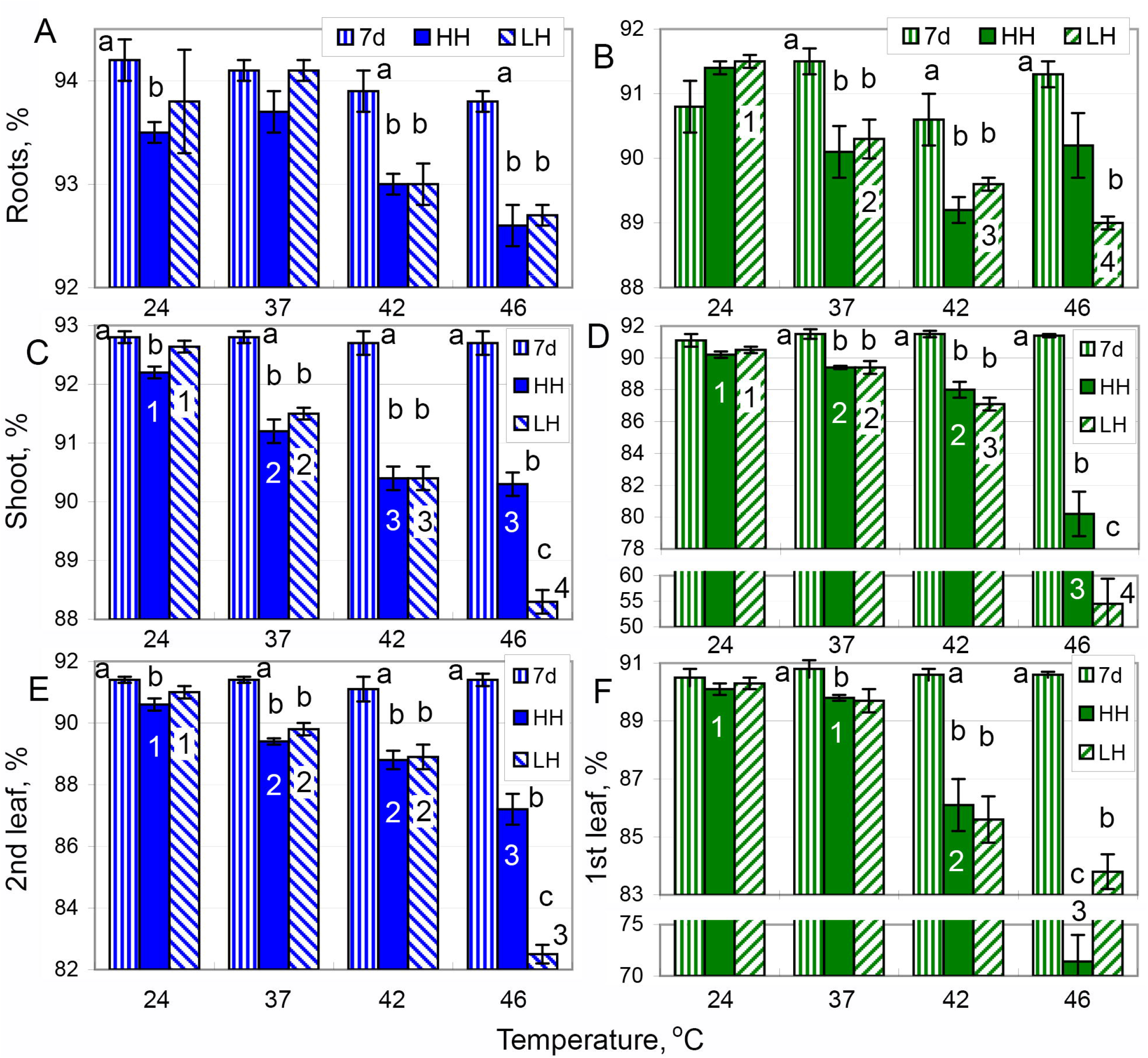
Effect of elevated temperature in contrast RH conditions on water content in organs of maize and barley plants. A, B – roots; C, D – shoot; E – second leaf, F – first leaf DW. A, C, E – maize; B, D, F – barley. All other designations are the same as in Fig. 1. The water content in all organs and complete statistical analysis are given in Supplementary Table S6.

The elevation of temperature stimulated water uptake by roots several times. In most cases, the water uptake remained at the same level under 37°C, 42°C, and 46°C; in maize plants grown at LH conditions, the elevation of temperature from 37°C to 46°C increased water uptake by about 40% (Fig 3, Supplementary Table S7). At the elevated temperatures, maize plants absorbed more water comparing with barley plants. It was general tendency; however, the difference was significant in two pairs of maize and barley plants grown at the same conditions (Fig 3, Supplementary Table S7). Under LH conditions, plants of the both species absorbed more water than plants grown in HH conditions. The difference was insignificant at the control temperature and in maize plants grown at 37°C; the growth of maize at 37°C was not inhibited at all (see above). Probably, the higher water uptake compensated the water loss through transpiration.

**Fig. 3.**
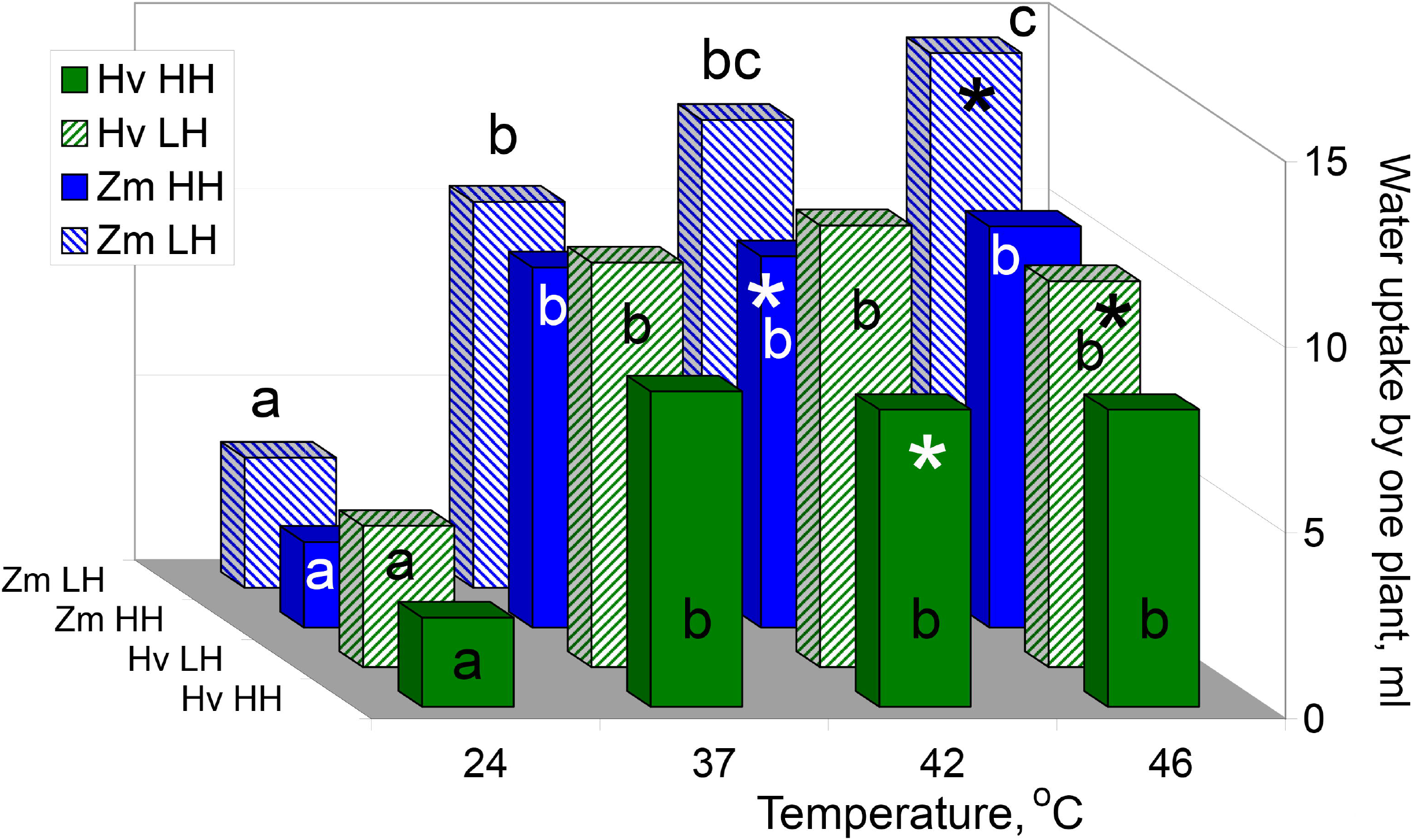
Water uptake by roots during 48 h at diverse regimes of temperature and air RH. The mean values are presented. Zm – *Zea mays*, Hv – *Hordeum vulgare*. a-c – differences between variants of temperature regime are significant at p ≤ 0.05. * - difference between two species in same conditions (°C, RH %) is significant at p ≤ 0.05. Partial statistical analysis is shown (see explanation at Fig. 1). The means ± SE data and complete statistical analysis are given in Supplementary Table S7.

### Chlorophyll content

The temperature 37°C stimulated small increase of photopigments in the both species. The content of Chl *b* was increased in the first leaves; Chls *a*, Chl *b*, and carotenoids were increased in the second leaves (Fig 4, Supplementary Table S8). Under the temperature 42°C, the levels of Chl *a* and Chl *b* were decreased in the both leaves of barley; in maize, the accumulation of photopigments was not changed. The effects of 37°C and 42°C did not depend on the air humidity (Fig 4, Supplementary Table S8).

**Fig. 4.**
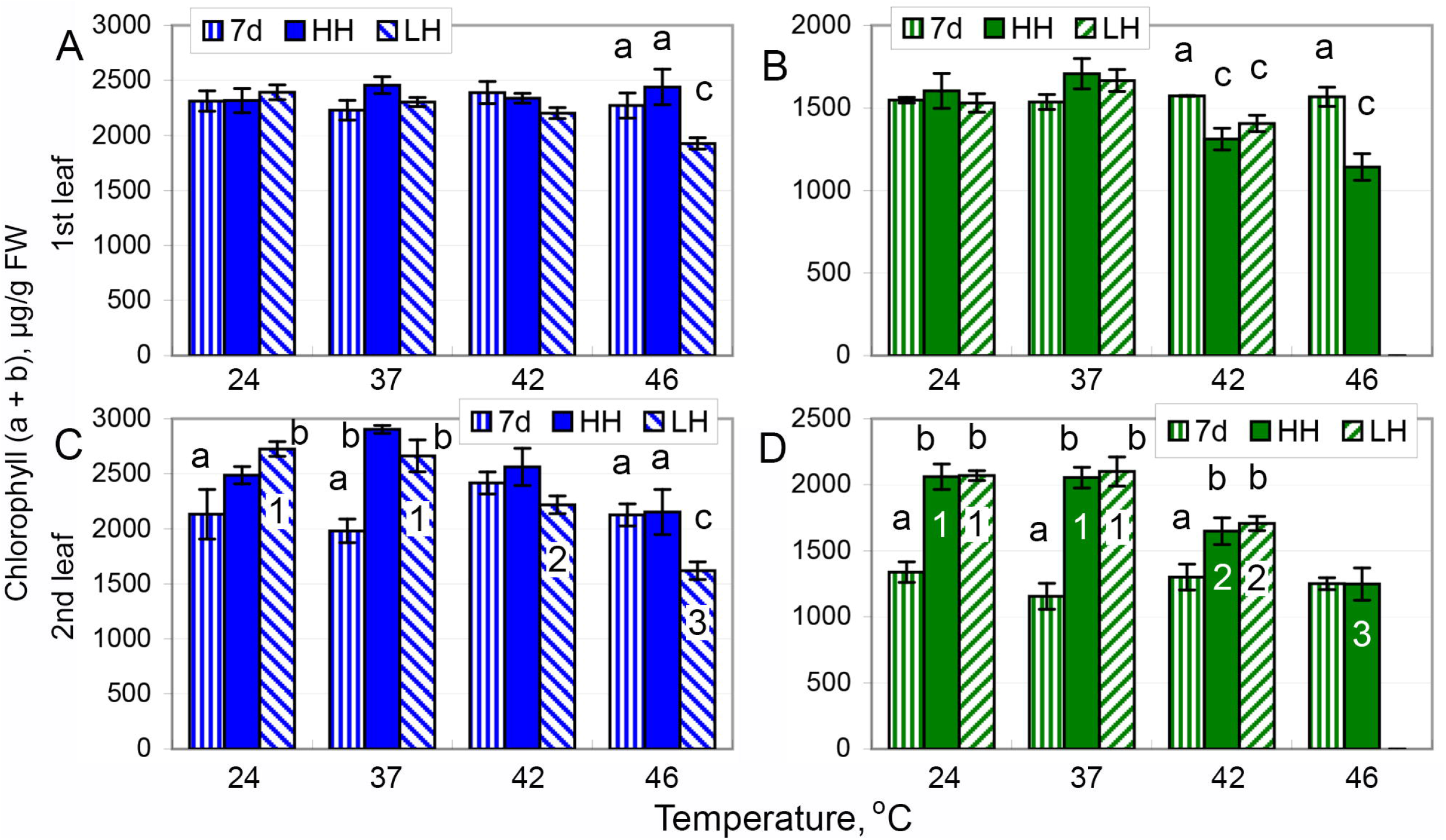
Effect of elevated temperature in contrast RH conditions on the content of Chl in the leaves of maize and barley. A, B – first leaf; C, D – second leaf. A, C – maize; B, D – barley. All other designations are the same as in Fig. 1. The comprehensive study of Chl *a*, Chl *b*, and carotenoids and complete statistical analysis are given in Supplementary Table S8.

Under the temperature 46°C, air humidity modified the impact of heat (Fig 4, Supplementary Table S8). In maize, the accumulation of photopigments was not changed at HH conditions; in LH conditions, the levels of Chls and carotenoids were decreased in the both leaves. In barley, the contents of Chls and carotenoids were decreased at HH conditions. Generally, the effect looks larger than under 42°C; the difference was significant for Chl *b* (Supplementary Table S8) and sum of Chls (Fig. 4D) in the second leaves. In LH conditions, the tips of barley leaves were damaged severely; the photopigments were not determined.

### Maximal Chl fluorescence and P_700_ light absorption in the dark

Minimal Chl fluorescence Fo was decreased at 37°C. The further elevation of temperature induced small extra decreases that were significant or not. The tendency from 37°C to 46°C was smooth in maize and nonlinear in barley. The differences between LH and HH variants were insignificant (Fig 5A, Supplementary Table S9).

**Fig. 5.**
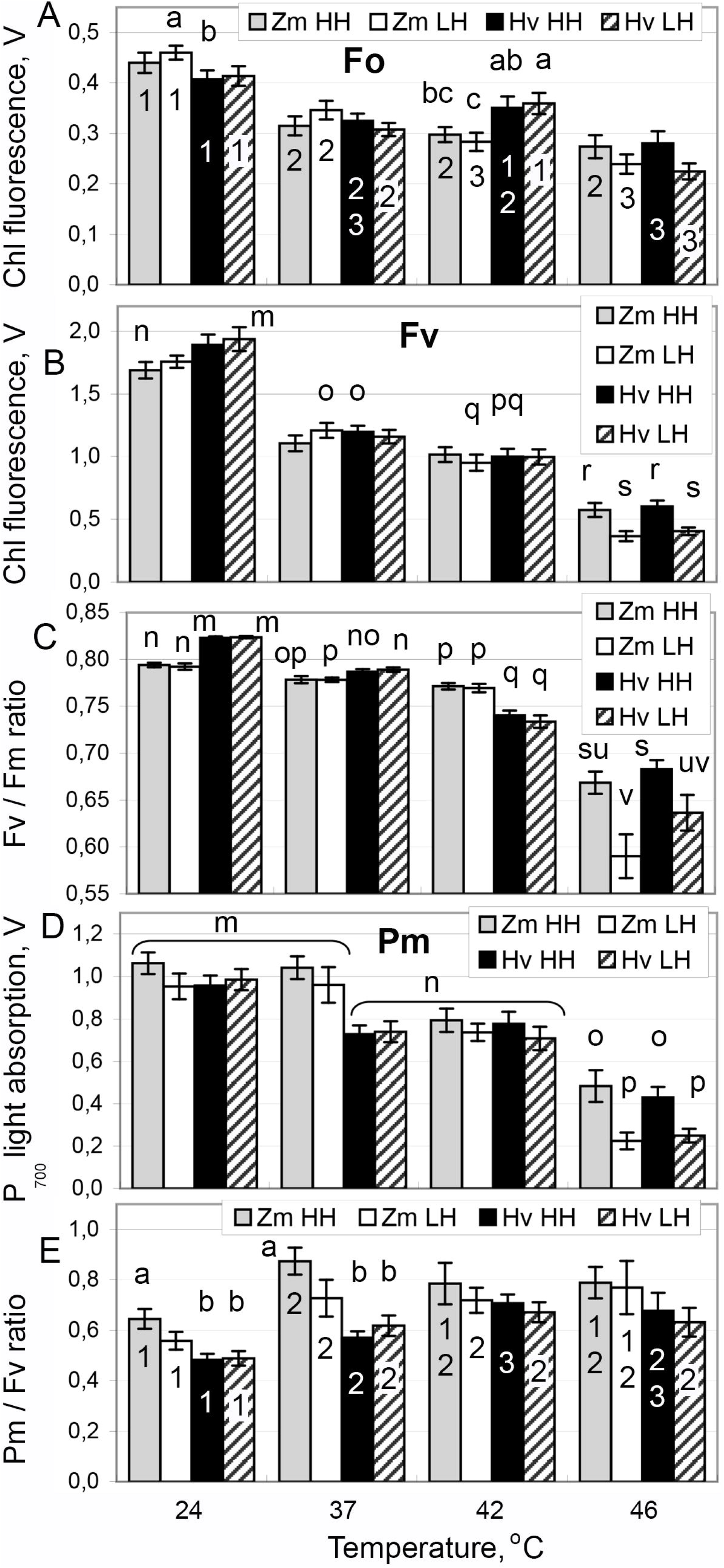
Basic photosynthetic values in the dark-adapted plants. A – Fo; B – Fv; C – Fv/Fm; D – Pm; E – Pm/Fv. Bars represent maize at HH (grey) and LH (white) conditions and barley at HH (black) and LH (hatched) conditions. In panels B-D, all the variants are compared to each other. m-v – differences between all variants (temperatre, air humidity, and species) are significant at p ≤ 0.05. All other designations are the same as in Fig. 1 and 3. The full dataset including Fm and complete statistical analysis are given in Supplementary Tables S9. The dynamics of Fv/Fm (C) are emphasized in Supplementary Fig. 1.

Maximal Chl fluorescence Fm and variable fluorescence Fv changed similarly. The changes of Fv are shown in Fig 5B; the changes of Fm can be found in Supplementary Table S9. The level of Fv decreased at 37°C and further decreased slightly at 42°C. At 46°C, Fv decreased to the very low level; in LH conditions, the fall was larger than in HH conditions.

The maximal quantum yield of PSII Fv/Fm was reduced gradually from 24°C to 42°C; in barley at HH conditions the tendency was extended to 46°C (Fig. 5C). In barley, Fv/Fm was higher in control and reduced faster with the temperature elevation comparing with maize (Supplementary Fig S1). At 46°C, Fv/Fm decreased drastically in three variants; in LH conditions, the fall was larger than in HH conditions (Fig 5C).

The maximal change of P_700_ light absorption Pm demonstrated qualitative difference between species. In barley, Pm felt at 37°C and remained at the same level at 42°C (Fig 5D) that resembles changes of Fv and Fo. In maize, Pm was not inhibited at 37°C; at 42°C Pm was reduced to the same level as in barley. At 46°C, Pm was decreased more in the both species; in LH conditions, the fall was larger than in HH conditions.

The value Pm reflects quantity of PSI reaction centers; Fv is attributed to the activity of all PSII reaction centers. The ratio Pm/Fv estimates balance between amounts of functionally active PSI and PSII (Lysenko *et al*., 2020). The elevation of temperature increased the value of Pm/Fv ratio (Fig 5E); the increase of this ratio suggests a shift of the balance in favor of PSI. In maize, Pm/Fv was increased at 37°C and remained at this level under higher temperatures. In barley, Pm/Fv increased gradually from 24°C to 42°C; the increase between 37°C and 42°C was significant in HH variant only (Fig 5E, Supplementary Table S9). Under the control temperature 24°C, the ratio Pm/Fv was higher in maize comparing with barley; at 42°C, Pm/Fv ratios were similar in the both species. The air humidity did not influence Pm/Fv ratio (Fig 5E).

### Photochemical activity of PSII

After the measurement in dark adapted state, the actual photosynthetic activity under illumination was recorded using IC method. During the *in vivo* experiment, plants were adapted to the light intensity 60-80 μmol photons m^−2^ s^−1^; therefore, AL intensity 70 μmol photons m^−2^ s^−1^ was chosen for IC. In PAM analysis, the photochemical activity of PSII is attributed to the difference between maximal and stationary Chl fluorescence in light: Fm′ – Fs = ΔF (Lichtenthaler *et al*., 2005). The photochemical activity of PSII can be analyzed with the use of three coefficients: qP = ΔF/Fv′, Φ_PSII_ = ΔF/Fm′, and X(II) = ΔF/Fv. The dynamics of qP and X(II) are shown in Fig. 6; dynamics of Φ_PSII_ is represented in Supplementary Fig. S2. All the three coefficients demonstrated the same general tendency and some coefficient specific features.

**Fig. 6.**
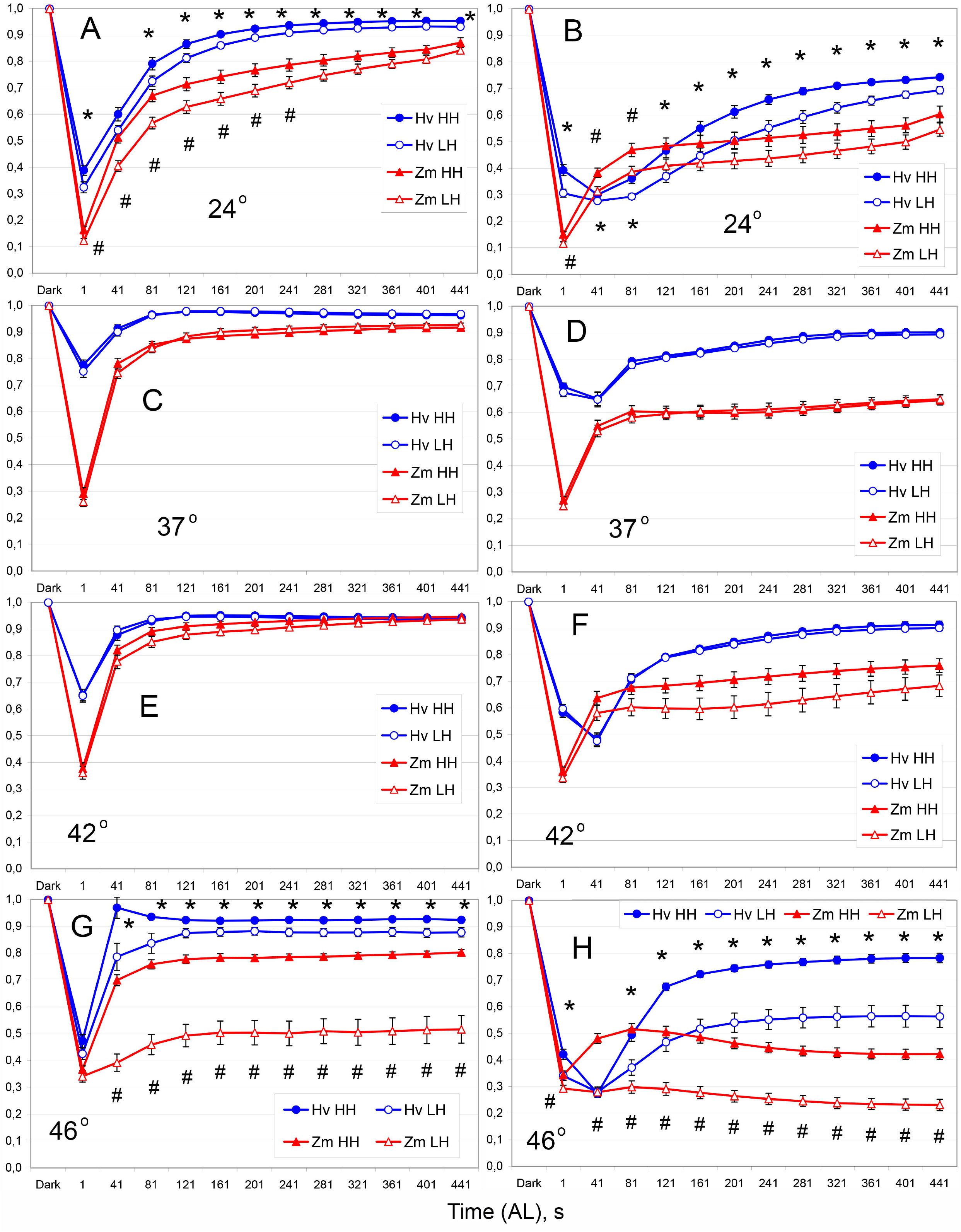
Dynamics of photochemical coefficients of PSII in IC at AL 70 μmol photons m^−2^ s^−1^. Effect of air humidity at the four distinct temperatures. A, C, E, G – qP; B, D, F, H – X(II) (dynamics of Φ_PSII_ are represented in Supplementary Fig. S2). A, B – 24°C; C, D – 37°C; E, F – 42°C; G, H – 46°C. Blue lines and circles represent barley (Hv); red lines and triangles represent maize (Zm). Filled symbols show HH conditions; open symbols show LH conditions. Data are presented as means ± SE. * - in barley, corresponding means at HH and LH conditions are different significantly at p ≤ 0.05. # - in maize, corresponding means at HH and LH conditions are different significantly at p ≤ 0.05. At 42°C (F), dynamics of X(II) (and φ_PSII_ also) at steady state phase are different significantly in maize (see Supplementary Commentary 1). The same data are presented in an alternative manner for analyzing temperature effect in the both species at both RH conditions (Supplementary Fig. S3).

In control (24°C), plants grown at HH exhibited the higher level of photochemical usage of light energy comparing with plants grown at LH conditions (Fig. 6A,B, Supplementary Fig. S2). The elevation of temperature to 37°C, stimulated the photochemical reactions of PSII; as the result, the difference between HH and LH conditions disappeared (Fig. 6C,D, Supplementary Fig. S2). At the temperature 42°C, all the photochemical coefficients remained higher than in control. In barley, air humidity did not affect photochemical processes at 42°C. In maize, the photochemical reactions were lowered in LH conditions. The difference between HH and LH variants was diminished in the row X(II) > Φ_PSII_ > qP (Fig. 6E,F, Supplementary Fig. S2). The pairwise comparison of the data points revealed no significant difference. The linear regression analysis demonstrated the significant difference between steady state phases (41-441 s of AL) of IC dynamics (Fig. 6F); in LH conditions, the dynamics of X(II) (p < 1e^−13^) and Φ_PSII_ (p = 2.1e^−5^) were lower than in HH conditions (Supplementary Commentary 1).

Under the temperature 46°C, the photochemical reactions of PSII were decreased below (maize) or mostly below (barley) the control level. Under nearly lethal HS, the effect of air humidity was obvious in the both species. In LH conditions, the fall of X(II), Φ_PSII_, and qP was much higher than in HH conditions (Fig. 6G,H, Supplementary Fig. S2). At 46°C, the differences between HH and LH were much larger than in the control except for qP in barley (Fig. 6, Supplementary Fig. S2).

The data obtained were organized to reveal any effect of air humidity (Fig. 6, Supplementary Fig. S2). To trace the effect of temperature, the same data were revisualized. In Supplementary Fig. S3, each panel shows all the temperature dependent curves (24-46°C) for the particular species at particular RH conditions. To prove the statements in the previous paragraph, the readers are referred to Supplementary Fig. S3.

The IC analysis was performed at the single light intensity. Reaction of photosynthetic apparatus may vary depending on AL intensities; experts in photobiology recommended to study activities of photosystems at various AL intensities (Lichtenthaler *et al*., 2005; Kalaji *et al*., 2014). Therefore, the investigation was extended with the application of RLC method that enables to study wide range of light intensity spectra. The RLC dynamics of qP and X(II) are shown in Fig. 7; dynamics of Φ_PSII_ is represented in Supplementary Fig. S4. For the analysis of temperature dependent effect, these data are revisualized in Supplementary Fig. S5.

**Fig. 7.**
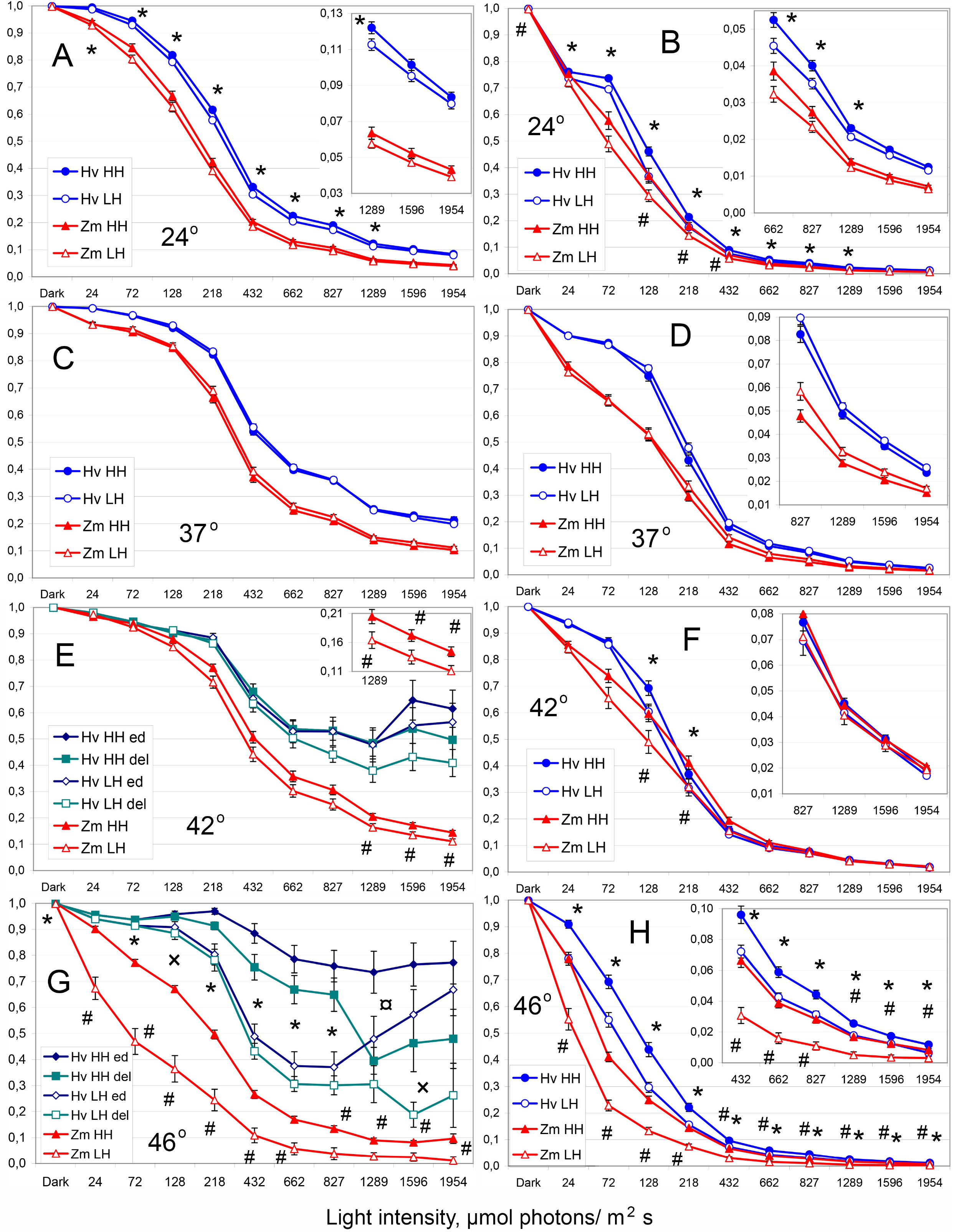
Dynamics of photochemical coefficients of PSII in RLC at diverse AL intensities. Effect of air humidity at the four distinct temperatures. A, C, E, G – qP; B, D, F, H – X(II) (dynamics of φ_PSII_ are represented in Supplementary Fig. S4). A, B – 24°C; C, D – 37°C; E, F – 42°C; G, H – 46°C. The insets show the corresponding small values with the higher resolution. In barley at 42°C (E) and 46°C (G), values Fo′>Fs caused incorrect qP values (see explanation in the text). The threshold level qP=1.05 was used; values qP>1.05 or negative (if Fo′>Fm′) were either deleted or edited to qP=1. Cyan lines and squares represent dynamics after deletion (del) of bad values; dark blue diamonds show dynamics after edition (ed) of bad values to qP=1. For E and G panels: * - corresponding means at HH and LH conditions are different significantly in the both cases (del and ed), p ≤ 0.05; × - difference between “del” variants is significant only, p ≤ 0.05; ¤ - difference between “ed” variants is significant only, p ≤ 0.05. All other designations are the same as in Fig. 6. The same data are presented in an alternative manner for analyzing temperature effect (Supplementary Fig. S5).

In control (24°C), barley plants grown at LH showed small decrease of photochemical coefficients comparing with counterparts grown at HH. Maize plants demonstrated similar tendency; due to higher variance, the only differences of X(II) at the AL intensities 128-432 μmol photons m^−2^ s^−1^ were significant. Similar to IC, the RLC dynamics of all the three photochemical coefficients were increased at 37°C (Supplementary Fig. S5); any differences between HH and LH variants were disappeared (Fig. 7, Supplementary Fig. S4). At the temperature 42°C, the photochemical activity of PSII remained higher than in control according to all the coefficients (Supplementary Fig. S5). In LH conditions, the photochemical coefficients were decreased comparing with HH conditions. The decrease was relatively small and revealed at the light intensities higher than 72 μmol photons m^−2^ s^−1^; the distribution of points with the significant differences varied between the coefficients qP, Φ_PSII_, and X(II) and overlapped partially (Fig. 7, Supplementary Fig. S4).

Under the nearly lethal HS (46°C), the photochemical reactions of PSII were decreased to the control level or below; in barley, qP level only remained higher than in control (Supplementary Fig. S5). At 46°C, the air humidity affected the activity of PSII much. The LH conditions stimulated larger decrease of all the photochemical coefficients comparing with HH conditions; the differences between HH and LH variants were obvious in the whole spectra of AL intensity (Fig. 7, Supplementary Fig. S4).

Barley plants kept high openness of PSII (qP) by means of nearly complete elimination of closed PSII (Fs – Fo′ ≈ 0). In such cases, Fo′-determination with far-red light induced values Fo′>Fs frequently and even Fo′>Fm′ rarely. The coefficient qP calculated from such data exceeded 1 (>100%) or was negative (<0%) correspondingly that is nonsense. These incorrect data cannot be exploited for the calculation of qP. First, these data points were discarded; however the elimination of data with minimal level of closeness decreased artificially the openness of PSII. Second, these data points were considered with no closed PSII (Fs – Fo′ = 0); therefore, they are open completely and qP=1. In this approach, openness of PSII will be increased artificially to a lesser extent. The both types of data edition are presented in Fig. 7E,G and Supplementary Fig S5. The calculation of X(II) and Φ_PSII_ were not influenced: Fm′ was larger than Fs always and Fo′ is not used for their calculation.

### Photochemical activity of PSI

The only coefficient Y(I) is used for the assessment of photochemical activity of PSI; therefore, the data of IC and RLC analyses were compared in parallel (Fig. 8, Supplementary Fig. S6). In control (24°C), the quantum yield Y(I) was lower in LH comparing with HH conditions. In IC curves measured at AL 70 μmol photons m^−2^ s^−1^, the both species demonstrated lower Y(I) levels in LH conditions (Fig. 8A). In RLC at higher AL intensities, barley demonstrated the decrease of Y(I) values in LH conditions also; in maize, the tendency was not obvious and significant at 432 μmol photons m^−2^ s^−1^ only (Fig. 8B).

**Fig. 8.**
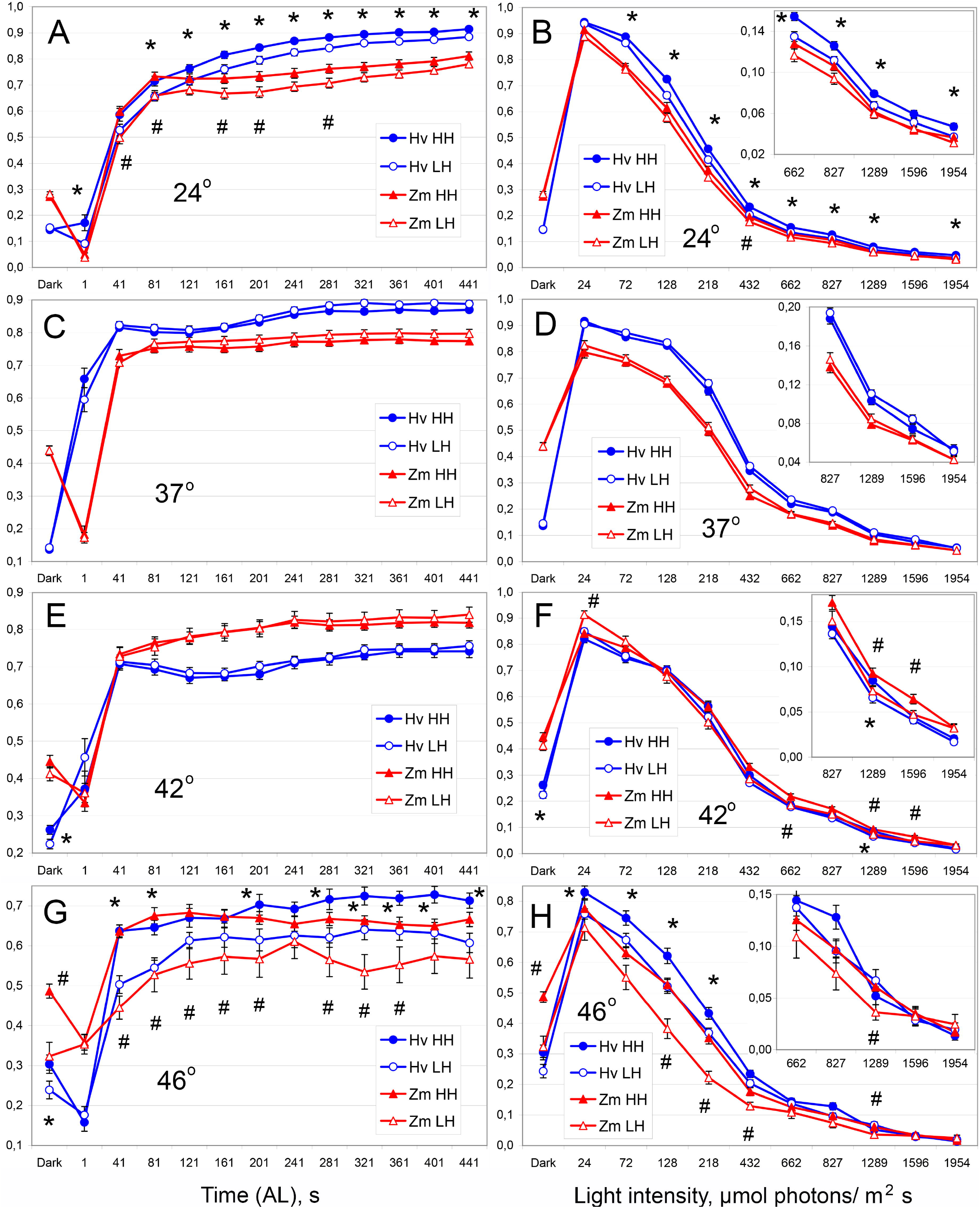
Dynamics of the quantum yield Y(I) of PSI. Effect of air humidity at the four distinct temperatures. A, C, E, G – IC (AL 70 μmol photons m^−2^ s^−1^); B, D, F, H – RLC. A, B – 24°C; C, D – 37°C; E, F −42°C; G, H – 46°C. The insets show the corresponding small values with the higher resolution. All designations are the same as in Fig. 6. The same data are presented in an alternative manner for analyzing temperature effect (Supplementary Fig. S6).

The elevation of temperature to 37°C stimulated PSI to achieve nearly the same steady state level in IC faster than in control (Supplementary Fig. S6). The early reaction demonstrated species specificity. The first second of illumination decreased Y(I) to zero level in maize and, vice versa, increased Y(I) closely to the steady state level in barley (Fig. 8C). The similar tendency was observed at 42°C; though, it was less obvious (Fig. 8E). In RLC at the higher AL intensities, temperature 37°C stimulated increase of Y(I) comparing with the control (Supplementary Fig. S6). The differences between HH and LH variants were absent at 37°C (Fig. 8C,D).

The further elevation of temperature influenced PSI diversely in maize and barley. In maize, the dynamics of Y(I) at 37°C and 42°C were similar in the both IC and RLC (Supplementary Fig. S6). In barley, the dynamics of Y(I) at 42°C were decreased below the level observed at 37°C; the decrease was observed in the both IC and RLC also (Supplementary Fig. S6). In IC, the steady state Y(I) level was reached faster (in 40 s) than in control; however; the steady state level itself was lower than in control (Fig. 8E, Supplementary Fig. S6). In RLC under the moderate AL intensities (200-800 μmol photons m^−2^ s^−1^), the values of Y(I) at 42°C were lower than at 37°C and higher than in control; at the highest AL intensities (1.5-2 mmol photons m^−2^ s^−1^), they were below the control level (Supplementary Fig. S6).

At 42°C, air RH conditions did not influence the Y(I) dynamics in IC at 70 μmol photons m^−2^ s^−1^ (Fig. 8E). Under higher AL intensities, LH conditions stimulated very small decrease of Y(I); the effect was significant in three points in maize and in a single point in barley (Fig. 8F).

**Fig. 9.**
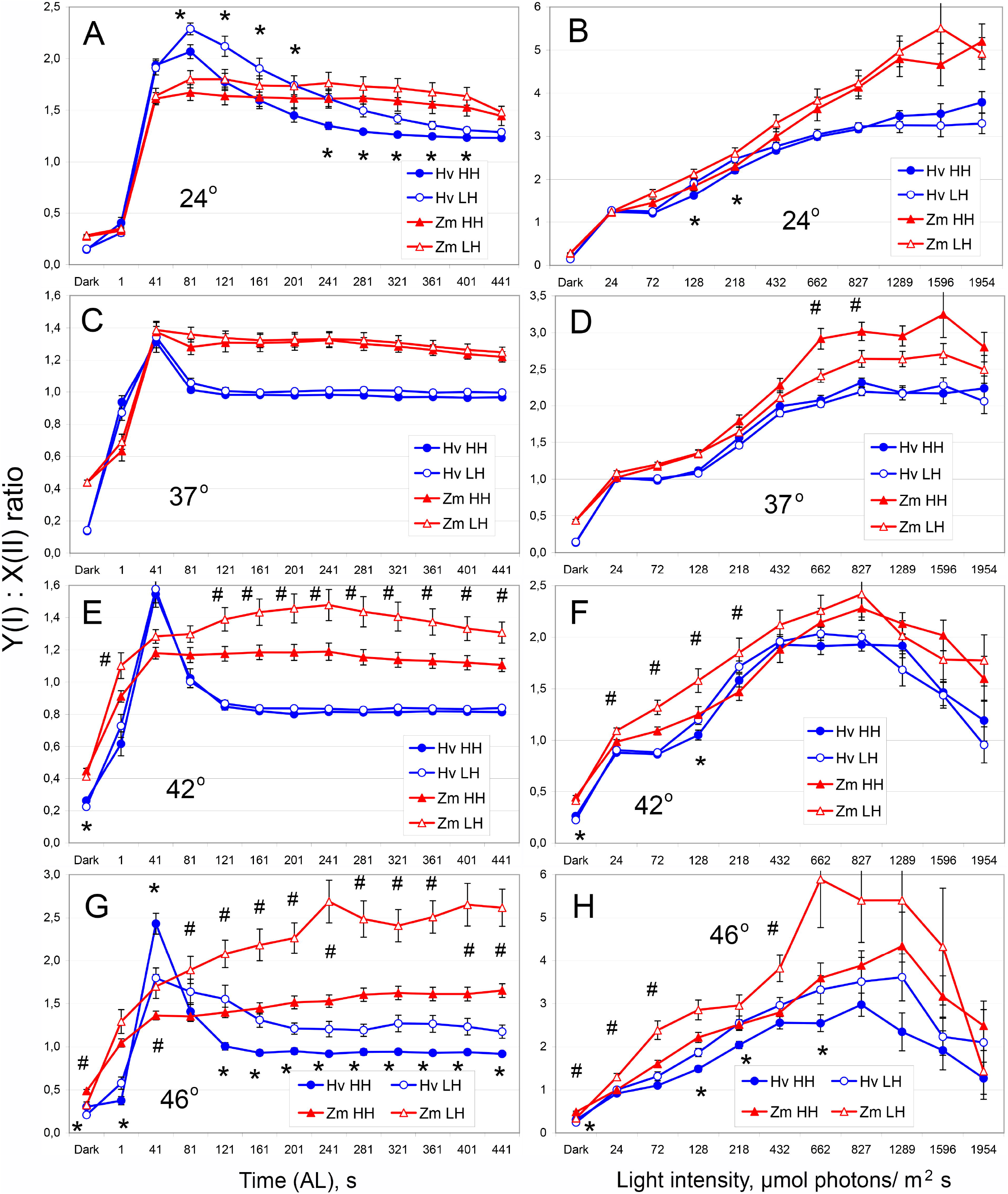
The balance between photochemical activities of PSI (Y(I)) and PSII (X(II)). Effect of air humidity at the four distinct temperatures. A, C, E, G – IC (AL 70 μmol photons m^−2^ s^−1^); B, D, F, H – RLC. A, B – 24°C; C, D – 37°C; E, F −42°C; G, H – 46°C. All designations are the same as in Fig. 6. The same data are presented in an alternative manner for analyzing temperature effect (Supplementary Fig. S7).

At the nearly lethal temperature 46°C, the dynamics of Y(I) felt below the control level under low light intensities (till 128 μmol photons m^−2^ s^−1^) in the both IC and RLC (Fig. 8G,H, Supplementary Fig. S6). At the moderate AL intensities, Y(I) was at the control level mainly; at the highest AL intensities, Y(I) decreased below control level once again (Fig. 8H, Supplementary Fig. S6). The LH conditions, stimulated larger decrease of Y(I) than HH conditions; it was observed in the both species and at AL intensities up to 1300 μmol photons m^−2^ s^−1^ (Fig. 8G,H). At 46°C, the LH dependent decrease of Y(I) was larger than in control (24°C); however, it looks smaller than the corresponding decrease of PSII photochemical activity at 46°C (Fig. 6, 7, Supplementary Fig. S2, S4).

### PSI vs PSII balance

The coefficients Y(I) and X(II) are calculated in analogous manner and can be compared to each other (Lysenko *et al*., 2020); therefore, the ratio Y(I)/X(II) can be used for estimating the balance between photochemical activities of PSI and PSII.

The ratio Y(I)/X(II) was calculated for IC for the first time and revealed qualitative difference between species. In the second leaves of maize, the dynamics of the ratio was monophasic. During the first 40 s of illumination, the ratio Y(I)/X(II) was increased and reached the steady state level (Fig. 9A,C,E) or turned to further slow growth (Fig. 9G). The first leaves of barley demonstrated biphasic dynamics. During the first 40 s of illumination, the ratio Y(I)/X(II) was increased and reached a peak; then the ratio turned to decrease and reached the steady state level. The steady state level was achieved fast (Fig. 9C) or slow (Fig. 9A). The values of ratio Y(I)/X(II) were higher in maize (C4 species) than in barley (C3 species) at any conditions of AL except the peak in barley (Fig.9; Supplementary Fig. S7).

The elevation of temperature to 37°C and 42°C decreased the ratio Y(I)/X(II), i.e. shifted the balance in favor of PSII; at nearly lethal temperature 46°C the ratio Y(I)/X(II) was returned to the control level in the most cases (Supplementary Fig. S7). In maize plants grown at 46°C and LH, the ratio Y(I)/X(II) increased above the control level that means shift of the balance in favor of PSI (Supplementary Fig. S7). Another one temperature dependent trait was found in RLC. The increase of AL intensity was followed with the increase of ratio Y(I)/X(II). At 24°C and 37°C, the increase was linear or reached a plateau (Fig 9B,D). At the higher temperatures 42°C and 46°C, the tendency achieved a peak at the light intensity about 800 μmol photons m^−2^ s^−1^; the further increase of light intensity caused the decrease of Y(I)/X(II) ratio (Fig. 9F,H).

The LH conditions increased the ratio Y(I)/X(II), i.e. shifted the balance in favor of PSI (Fig. 9). In control, the LH induced increase was significant in barley at low AL intensities (Fig. 9A,B); in maize, it was smaller and insignificant. At the temperature 37°C, the difference was disappeared (Fig. 9C,D). In maize, an opposite tendency was detected at the higher AL intensities; the differences in two points were significant (Fig. 9D). Probably, this difference was accidental. At the temperature 42°C, the LH conditions increased the ratio Y(I)/X(II) in maize at the lower AL intensities (24-218 μmol photons m^−2^ s^−1^) (Fig. 9E,F). At the nearly lethal temperature 46°C, the ratio Y(I)/X(II) was increased in the both species; the increase was observed until the moderate light intensity (432-662 μmol photons m^−2^ s^−1^). In maize, the increase at 46°C was larger than at 42°C (Fig. 9G,H).

## Discussion

### Influence of HS and air RH on plant growth

The current analysis comprised three elevated temperatures. The temperature 37°C is considered as a moderate heat stress (Sharkey, 2005). This temperature slowed down the growth of maize and barley slightly. Maize plants accumulated less amount of mass in roots (Fig. 1A) and water in shoots (Fig. 2C); the levels of Chls and carotenoids were increased in the second leaves of HH variants (Fig. 4C, Supplementary Table S8), probably, due to water loss. Barley plants are more heat sensitive; therefore, they demonstrated more growth retardation. The growth of second leaves was slowed down according all the parameters studied; the accumulation of FW with stems was arrested completely. Barley whole shoots were delayed in the linear growth and accumulation of FW that was significant in HH variants (Fig. 1; Supplementary Tables S1-S5). The water content was decreased in the most of organs (Fig. 2, Supplementary Table S6). The content of Chl *b* was increased in the first leaves yet (Supplementary Table S8). In the current experiment, conditions of 37°C should be considered as mild (maize) or moderate (barley) HS irrespective of air humidity.

The temperature 42°C inhibited plant growth to larger extent. According to some parameters, the growth was abolished; e.g., FW of maize roots and DW of barley roots were not increased at all during 48 h of growth (Fig. 1A, Supplementary Table S1). In few cases, parameters were reduced after 48 h of growth at 42°C; e.g., FW of barley roots and width of maize leaves were smaller in 9-day old plants than in 7-day old plants taken into the experiment (Fig. 1B,I, Supplementary Table S4). The water content decreased in all organs of the both species. Contents of the both Chl *a* and Chl *b* were diminished in the second leaves of maize and both leaves of barley (Fig. 4, Supplementary Table S8). The small impact of air RH was revealed at 42°C in limited number of cases. In barley stems, the water content decreased more in LH conditions than in HH conditions (Supplementary Table S6). In the second leaves of maize, the content of Chl *b* decreased in LH conditions only (Supplementary Table S8). The current conditions of 42°C can be considered as genuine HS with limited impact of air RH.

The temperature above 45°C can be lethal, probably, because of irreversible inactivation of PSII (Sharkey, 2005). Such high temperatures are generally applied for minutes (see Introduction) or few hours (Camejo *et al*., 2005; Yan *et al*., 2013). In the current experimental model, the each temperature was applied for 48 h. To overcome possible lethality, we used continuous illumination that increased resistance to HS (Havaux and Strasser, 1990; Havaux *et al*., 1991; Kalituho *et al*., 2003). Poacea species own large potential for resistance to many abiotic stresses also. Plants survived 48 h at 46°C with active PSII and PSI that implies implementation of long term adaptive mechanisms.

The temperature 46°C reduced severely the plant growth in the both species. The growth was arrested completely according to many parameters of size and weight. Moreover, the loss of size and weight was revealed in many cases; after two days in the thermostat chambers, the size and/or weight parameters were smaller than in 7-day old plants before the experiment (Supplementary Tables S1-S5). Surprisingly, the shoots of both species continued accumulating of DW at 46°C. The second leaves of the both species, the first leaves of barley, and the stems of maize at HH conditions demonstrated stable accumulation of DW at 46°C (Supplementary Tables S2-S5). Contrary, the roots lost DW at 46°C (Supplementary Table S1). The water content decreased in all organs of the both species; the effect was large in many cases (Fig. 2, Supplementary Table S6). The contents of Chl *a*, and Chl *b*, and Car were reduced (Fig. 4, Supplementary Tables S8). Frequently, the levels of all photosynthetic pigments were below their levels in 7-day old plants; it was revealed in the both leaves of maize at LH conditions and in first leaves of barley at HH conditions (did not measured at LH conditions).

The specific feature of 46°C treatment was reduction of organs by several times. The drastic losses of FW, DW, and water were found in barley mainly (Supplementary Tables S1-S5). In maize, the only example was found: root FW was decreased nearly twice (Fig. 1A).

At 46°C, the retardation effect depended on the air RH usually. In the both species, the degree of reduction of DW, FW, water content, and contents of Chls and carotenoids depended on the air RH conditions. At LH conditions, the inhibitory action was always more severe with the only exception. In the first leaves of barley, water content (%) was decreased more in HH conditions than in LH conditions (Fig. 2F). In this case, the balance between water and dry matter was governed with the great and diverse changes of dry matter. In HH conditions, DW was increased nearly twice (Fig. 1B) that reduced proportion (%) of water. In LH conditions, DW was decreased three times (Fig. 1B) that increased proportion (%) of water. Barley plants survived at 46°C in HH conditions; in LH conditions they were depressed greatly and died when the temperature treatment exceeded 48 h for extra 3-5 h. Therefore, some parameters were not measured in this variant. The current conditions of 46°C can be considered as nearly lethal HS with the large impact of air RH.

Plants were supplied with the unlimited source of a water based mineral media. The temperature elevation to 37°C stimulated the water uptake by roots several times (Fig. 3). Plants demonstrated no or minimal water loss from their organs including the leaves (Fig. 2, Supplementary Table S6). Probably, all (increased) transpiration was compensated with the increased water uptake. Under the higher temperatures 42°C and 46°C, the water uptake by roots was not increased further in most cases (Fig. 3, Supplementary Table S7). At 42°C and 46°C, plants lost more water; the large water loss was specific to leaves and greater in LH conditions than in HH conditions (Fig. 2, Supplementary Table S6). In barley, the water uptake remained at the same level at 37-46°C; however, it was always higher in LH than in HH conditions (Supplementary Table S7). Probably, barley exhausted all possibilities for increasing water uptake at 37°C. In maize at HH conditions, the temperature increase from 37°C to 46°C did not increase water uptake also (Fig. 3); the water loss from tissues was quite moderate (Fig. 2, Supplementary Table S6). The 37°C was mild stress for maize (see above); at 37°C the water uptake at the HH and LH conditions was equal (Supplementary Table S7). In LH conditions, maize plants increased water uptake along with the further temperature elevation (Fig. 3, Supplementary Table S7). The increased water uptake by roots did not compensate completely the water loss from leaves (due to transpiration, obviously). However, the water loss in maize was smaller than in barley (Fig. 2, Supplementary Table S6) and maize plants survived conditions of 46°C at LH for 48 h with no signs of severe damages.

### Influence of HS on PSII and PSI

The current research aimed to the effect of air RH on photosynthesis. The experimental model is of unusual type (see Introduction). It allows studying long term adaptation of photosynthetic apparatus to temperatures 42°C and even 46°C. Therefore, the effects of elevated temperatures themselves will be considered before the effect of air humidity.

Values measured in dark adapted plants shows the maximally possible signal and may be referred to an amount of photosynthetically active reaction centers. The maximal P_700_ light absorption in the dark (Pm) reflects an amount of active reaction centers of PSI (Dual-PAM_1e, 2009). The variable fluorescence in the dark (Fv) is referred to total amount of functionally active reaction centers of PSII; the minimal fluorescence in the dark (Fo) is attributed to processes in antenna complexes of PSII with small impact of antenna complexes of PSI (Lichtenthaler *et al*., 2005; Kalaji *et al*., 2014). The “antennal” minimal fluorescence Fo is proportional directly to their quantity and inversely to a connectivity between and inside antenna complexes.

The review (Allakhverdiev *et al*., 2008) stated that HS raises Fo level; the review referred readers to experimental works with short term (5-20 min) application of HS. After two days of HS, the level of Fo was lowered (Fig. 5A). The temperature elevation to 37°C decreased Fo. The further increase to 42°C imposed small additional effect: decrease (maize) or increase (barley). The elevation to 46°C resulted in the lowest Fo level (Fig. 5A, Supplementary Table S9). Probably, short term HS increased Fo through a disconnection of energy transfer in antenna complexes; during the long term exposure, plants improved connectivity. In barley, the rise of Fo at 42°C can be caused with disconnection of antenna complexes. Under the temperature 46°C, the reduction of Fo can be enhanced with the decrease of Chl content (Fig. 4) that suggests a reduction of antenna complexes.

The level of Fv was changed similar to Fo, however, with the larger amplitude. The temperature elevation to 37°C decreased Fv substantially. Further elevation to 42°C caused small extra decrease of Fv; elevation to 46°C was followed with another substantial decrease (Fig 5B, Supplementary Table S9). The Fv/Fm ratio decreased gradually along with the temperature elevation (Fig. 5C). It is well known that Fv/Fm is higher in C3 species and lower in C4 species (Kalaji *et al*., 2014). In control conditions, Fv/Fm was higher in barley (C3 plant) than in maize (C4 plant). Maize is more heat resistant species than barley. Therefore, tolerated HS decreased Fv/Fm in maize slowly than in barley; at 42°C, values Fv/Fm were higher in maize than in barley (Fig. 5C, Supplementary Fig. S1). Nearly lethal HS (46°C) decreased Fv/Fm drastically in the both species.

In contrast to PSII dependent parameters, Pm was unchanged at 37°C in maize; the decrease was observed at 42°C only. In barley, Pm was decreased at 37°C with no further decrease at 42°C. Levels of Pm were indistinguishable in the both species at 24°C and 42°C (Fig. 5D). Under the temperature 46°C, the next decrease of Pm was revealed that was similar in the both species.

The changes of Fv and Pm can be extrapolated to quantitative changes of PSII and PSI. The ratio Pm/Fv allows estimating a balance between reaction centers of PSII and PSI. The temperature elevation increased Pm/Fv ratio; therefore, the balance was shifted in favor of Pm value and PSI quantity. In maize, the ratio Pm/Fv was increased at 37°C and kept at this level at the higher temperatures. In barley, the increase was slower and reached high level at 42°C (Fig. 5E). In control, the ratio Pm/Fv was higher in maize; at 42°C the levels of Pm/Fv were similar in the both species. Under near lethal temperature (46°C), the absolute values were small that increased the variances.

All these data are in good agreement with the general view that PSI is more tolerant to HS than PSII (Allakhverdiev *et al*., 2008).

The actual photochemical activity is tested in the presence of AL. The photochemical activity of PSII is measured as difference Fm′ – Fs = ΔF (Lichtenthaler *et al*., 2005). Several equations use ΔF value for the estimation of PSII activity under light conditions. Photochemical coefficient qP = ΔF/Fv′ shows proportion of open PSII (ΔF) compared to all excited PSII both open and closed (Fv′) (Kalaji *et al*., 2014). Actual quantum yield of PSII Φ_PSII_ = ΔF/Fm′ shows proportion of photochemical quenching of energy (ΔF) compared to all energy in PSII reaction centers and antenna complexes (Fm′). This coefficient is best for the comparison with Fv/Fm ratio. Relative quantum yield of PSII X(II) = ΔF/Fv shows proportion of actual photochemical quenching in light (ΔF) compared to maximally possible photochemical quenching in dark (Fv). This coefficient is best for the comparison with the quantum yield of PSI Y(I) (Lysenko *et al*., 2020). The coefficient X(II) normalizes ΔF dynamics to a constant Fv value measured in darkness; therefore, dynamics X(II) reflects the dynamics of ΔF itself (Lysenko *et al*., 2020). Coefficients qP and Φ_PSII_ normalize ΔF to the values Fv′ and Fm′. They are measured simultaneously with ΔF and includes components ΔC = Fs - Fo′ and Fo′; ΔC and Fo′ have their own dynamics in light conditions (Lysenko, 2021). Therefore, the coefficients qP and Φ_PSII_ reflects both dynamics of ΔF and its balance with dynamics of ΔC and Fo′ in current light conditions. Therefore, dynamics of X(II), qP, and Φ_PSII_ are different despite they used the same numerator.

Surprisingly, the temperature elevation to 37°C and 42°C increased photochemical activity of PSII. It was observed in the both IC (Supplementary Fig. S3) and RLC (Supplementary Fig. S5). It was obvious according to all the coefficients. In barley at 42°C, Φ_PSII_ only was at the level of control or below it. Probably, this decline was caused with the dynamics of Fo′ because Fo′-independent coefficients qP and X(II) did not show such decreases (Supplementary Fig. S3, S5). Under nearly lethal HS (46°C), the photochemical activity of PSII was at the control level or reduced below it. In barley plants at high AL intensities, photochemical coefficient qP was higher at 46°C than in control (Supplementary Fig. S5). Barley plants reduced portion of closed PSII greatly. The decrease of closed PSII increased proportion of opened PSII.

One second of illumination decreased photochemical ability of PSII in the control plants (Fig. 6, Supplementary Fig. S3); then the photosynthetic apparatus was readapted to light and the photochemical ability restored few minutes later. Plants grown at elevated temperature acquired an ability to minimize this fall. In barley plants grown at 37°C, the fall at first second was reduced greatly; the steady state level was achieved faster also. The further elevation of temperature inhibited this ability. At 42°C, the fall was of intermediate size; at 46°C, the fall was returned back to the control level. In maize, similar tendency was observed; however, its amplitude was much smaller than in barley.

Simultaneously, one second of illumination increased greatly the quantum yield of PSI in barley plants grown at 37°C (Fig. 8, Supplementary Fig. S6. The further elevation of temperature inhibited this ability also. The effect was smaller at 42°C and disappeared at 46°C. In IC, plants grown at 37°C and 42°C reached steady state level of Y(I) faster than in control. In contrast to PSII activity, the steady state level of Y(I) was not increased. The steady state level of Y(I) remained at the control level in the both species at 37°C and in maize at 42°C; in barley, it was decreased at 42°C (Supplementary Fig. S6). At higher AL intensities in RLC, the temperature 37°C increased Y(I) above the control level; in maize, 42°C increased Y(I) also. In barley, Y(I) level at 42°C fluctuated around the control level in RLC. Under nearly lethal HS (46°C), the steady state level of Y(I) was decreased below the control level at 72 μmol photons m^−2^ s^−1^; at the higher AL intensities in RLC, Y(I) was at the control level or below it (Supplementary Fig. S6).

These data provide an unexpected conclusion: an elevated temperature stimulates photochemical processes in thylakoid membranes. The moderate elevation of temperature (37°C) increased activities of PSII and PSI. The further increase of temperature diminished this activation. The data obtained after 1 s of illumination suggests improving of electron transport between PSII and PSI at moderately elevated temperature. It can be explained by several factors. First, quantities of PSII and PSI were decreased, probably (Fig. 5). A reduced density of photosystems can ease traffic of electrons between them. Second, an elevation of temperature increases a fluidity of thylakoid membranes (Horvath *et al*., 1998). We kept the temperature(s) of treatment during the adaptation to darkness and PAM measurement to save any features of thermally treated plants including membrane fluidity. The physical state of lipid phase of thylakoid membranes influences processes associated with these membranes (Goss *et al*., 2005). A moderate increase of fluidity can stimulate transport of electron shuttles along the membrane also. In 7-day old barley plants, an elevation of temperature (40°C, 3 h) stimulated the reoxidation of plastoquinone pool (Pshybytko *et al*., 2008). This can be caused by an increased mobility of plastoquinones in more fluid membranes. An elevated temperature stimulated electron exchange between plastoquinones in membrane and molecules in stroma (reviewed by Sharkey, 2005). At last, plants were able to synthesize new proteins or other molecules during 48 h adaptation to HS (Horvath *et al*., 1998; Tanaka *et al*., 2000; Sharkey, 2005; Allakhverdiev *et al*., 2008). Therefore, a temperature elevation can benefit some photochemical processes; under the higher temperatures, harmful effects can overcome a beneficial impact.

In darkness, the quantum yield Y(I) is larger in C4-plant maize comparing with C3-plant barley (Fig. 8). This suggests higher impact of cyclic electron transport or other mechanism of electron donation from stromal reductants to PSI in maize.

Relative activities of the PSI and PSII were compared with the use of ratio Y(I)/X(II) (Lysenko *et al*., 2020). The IC demonstrated differences between two species in the first min of AL conditions. In the second leaves of maize, the both X(II) and Y(I) demonstrated similar dynamics (Fig. 7, 8, Supplementary Fig. S3, S6); therefore, the dynamics of Y(I)/X(II) was simple and comprised fast raise with subsequent plateau or slow changes (Fig. 9, Supplementary Fig. S7). In the first leaves of barley, PSI and PSII adapted their photochemical abilities to light conditions with the different rates. After 40 s in light conditions, quantum yield of PSI Y(I) reached a high level (Supplementary Fig. S6) whereas relative quantum yield of PSII X(II) remained at the lowest level (Supplementary Fig. S3); in this time, Y(I)/X(II) ratio reached a peak (Supplementary Fig. S7). Further on, PSII photochemical activity was increased that decreased the ratio Y(I)/X(II) to a steady state level. This data showed that photochemical activity of PSI may be independent partially from PSII activity. Probably, barley first leaves had a reach pool of stromal reductants that were able supporting PSI activity for about 1 min through the cyclic electron transport or other pathways (Havaux, 1996; Yamane *et al*., 2000).

In barley, the steady state Y(I)/X(II) level was about 1.25 in control and decreased to 0.8-1 at 37°C and 42°C (Fig. 9, Supplementary Fig. S7). This means that PSI was slightly more active in control whereas PSII was slightly more active at tolerated HS. In maize, the steady state level of Y(I)/X(II) was higher; in control, it was around 1.5 and decreased to about 1.25 at 37°C and 42°C (Supplementary Fig. S7, Fig. 9). In maize, PSI was more active than PSII; tolerated HS shifted the balance in favor of PSII also. Considering maximal activity in dark adapted state as a quantitative measure of photosystems, we can suggest that tolerated HS shifted the quantitative balance in favor of PSI (Fig. 5E); such “better” stoichiometry may underlay increase of PSII relative activity. Under the near lethal HS (46°C), Y(I)/X(II) level remained at the decreased level (barley HH), turned back to the control level (barley LH, maize HH), or increased above the control level (maize LH); the last case means shift of the relative activities in favor of PSI comparing with the control level (Supplementary Fig. S7).

We notice that the steady state levels of the ratio Y(I)/X(II) were close to 1 in most cases; this means near equal activities of the both photosystems. Therefore, the estimation looked valid.

The effect of temperature elevation was similar when the ratio was studied in a wide range of AL intensities with RLC method. At the temperatures 37°C and 42°C, the ratio Y(I)/X(II) was decreased; i.e., it was shifted in favor of PSII. At 46°C it was reversed back to the control level at low and moderate light intensities (Supplementary Fig. S7). At the temperatures 24°C and 37°C, the increase of AL intensity enlarged the ratio Y(I)/X(II) continuously or it reached a plateau (Fig. 9B,D). At higher temperatures, the curves showed a critical point where the tendency was reversed. At 42°C and 46°C, the ratio Y(I)/X(II) raised until about 1000 μmol photons m^−2^ s^−1^; next the increase of AL intensity decreased the ratio Y(I)/X(II) (Fig. 9F,H). Activity of PSII decreased faster at earlier stages of RLC and declined to a very low level. When the electron donation from PSII, cyclic electron flow and/or stromal reductants was exhausted then activity of PSI turned to be reduced faster. The feature was universal and revealed in the both species at both conditions of air RH (Fig. 9, Supplementary Fig. S7).

### Influence of air RH on PSII and PSI

The conditions of air RH influenced activities of photosystems in control. The photochemical activity of PSII was decreased under LH conditions comparing with HH conditions. In IC at 72 μmol photons m^−2^ s^−1^, it was revealed in the both species (Fig. 6A,B, Supplementary Fig. S2). Under wide spectrum of AL intensities, it was revealed in barley according to all the coefficients (Fig. 7A,B, Supplementary Fig. S4). In maize, the coefficients qP and Φ_PSII_ demonstrated no significant differences (Fig. 7A, Supplementary Fig. S4). According to X(II), the difference at 72 μmol photons m^−2^ s^−1^ was insignificant; at the next three light intensities 128-432 μmol photons m^−2^ s^−1^, the differences were significant (Fig. 7B). The coefficient X(II) reflects dynamics of ΔF only; in the coefficients qP and Φ_PSII_ dynamics of ΔF is influenced with simultaneous alterations of (Fs-Fo′) and Fo′ (discussed above).

The quantum yield of PSI was decreased in a similar way. In control, barley plants demonstrated the decrease of Y(I) in LH conditions comparing with HH conditions; it was revealed in the both IC and RLC (Fig. 8A,B). In maize, LH conditions decreased Y(I) level in IC (Fig. 8A); in RLC, the difference was negligible and significant at 432 μmol photons m^−2^ s^−1^ only (Fig. 8B).

Under moderate HS (37°C), plants increased the photochemical activities of PSII and PSI. At 37°C, air RH did not influence the activities of PSII and PSI (Fig. 6C,D, Supplementary Fig. S2, Fig. 7C,D, Supplementary Fig. S4, Fig. 8C,D); the effect of LH conditions was canceled.

Under genuine HS (42°C), LH conditions induced small decrease of PSII and PSI activities once again. IC measured at 72 μmol photons m^−2^ s^−1^ demonstrated minimal differences between HH and LH conditions. The air RH did not influence the photochemical activity of PSI in the both species (Fig. 8E) and PSII in barley (Fig. 6E,F, Supplementary Fig. S2). In maize, LH conditions decreased the photochemical activity of PSII. The steady state levels of X(II) (Fig. 6F) and Φ_PSII_ (Supplementary Fig. S2) were lower in LH comparing with HH conditions; the difference was small but significant according to the linear regression analysis (Supplementary Commentary 1). The coefficient qP showed no differences between HH and LH conditions (Fig. 6E); consequently, the LH conditions decreased a number of the both open and closed PSII proportionally in maize.

In wide spectrum of AL intensity, the effect of air RH at 42°C was more extensive The LH conditions decreased the photochemical activity of PSII in the both species. The inhibitory effect was revealed at AL intensities higher than the illumination level during the experiment (60-80 μmol photons m^−2^ s^−1^). According to X(II), LH decreased PSII activity at 128-218 μmol photons m^−2^ s^−1^ in the both species (Fig. 7F). According to Φ_PSII_, the difference was appeared at 128-218 μmol photons m^−2^ s^−1^ and continued to 662 μmol photons m^−2^ s^−1^ (maize) or 1596 μmol photons m^−2^ s^−1^ (barley) (Supplementary Fig. S4). In maize, the coefficient qP was lowered in LH conditions from 128-218 μmol photons m^−2^ s^−1^ also; however, the decrease reached significant level at the highest AL intensities only (>1 mmol photons m^−2^ s^−1^) (Fig. 7E). In maize, LH conditions decreased PSI activity at the moderate and high AL intensities also; the differences were significant at 662, 1289, and 1596 μmol photons m^−2^ s^−1^ (Fig 8F). In barley, PSI was not influenced with air RH conditions; significant difference at a single point should be considered as a chance effect (Fig 8F). The most of these differences were small.

Under nearly lethal HS (46°C), LH conditions imposed large inhibitory effects on the both photosystems. The maximal values in dark adapted state Fv and Pm were decreased in LH comparing with HH variants (Fig. 5B,D, Supplementary Table 9); this suggests that LH conditions reduced amounts of active PSII and PSI reaction centers. Their relative photochemical activities were reduced in LH conditions also. Activity of PSII was reduced in the both species; it was revealed with the both IC and RLC methods (Fig. 6G,H, Supplementary Fig. S2, Fig. 7G,H, Supplementary Fig. S4). Activity of PSI was reduced in the both species and in the both IC and RLC also (Fig. 8G,H). In RLC, LH conditions decreased Y(I) significantly at low AL intensities in barley and moderate and high AL intensities in maize (Fig. 8H).

The ratio of maximal values Pm/Fv was nod influenced with RH conditions (Fig. 5E). This suggests that amounts of active PSI and PSII were decreased proportionally. The LH conditions shifted the balance of relative activities in favor of PSI. In barley, the ratio Y(I)/X(II) was increased at the control temperature 24°C. This effect was revealed at the relatively low light conditions: at AL 72 μmol photons m^−2^ s^−1^ in IC (Fig. 9A) and at 128-218 μmol photons m^−2^ s^−1^ in RLC (Fig. 9B). Similar small shift can be recognized in IC of maize (Fig. 9A); it was significant according to a nonparametric statistic (Supplementary Commentary 2). Under moderate HS (37°C), RH conditions did not influence Y(I)/X(II) ratio (Fig. 9C,D); in maize, the significant differences at AL 662 and 827 μmol photons m^−2^ s^−1^ should be referred to an effect by chance (Fig. 9D). Under genuine HS (42°C), LH conditions increased the ratio Y(I)/X(II) at relatively low light conditions also. In maize, the ratio was increased at 72 μmol photons m^−2^ s^−1^ in IC (Fig. 9E) and at AL 24-218 μmol photons m^−2^ s^−1^ in RLC (Fig. 9F). In barley, the increase was found in RLC only. Similar to the control, the ratio was increased at 128-218 μmol photons m^−2^ s^−1^ (Fig. 9F); the difference was significant at 128 μmol photons m^−2^ s^−1^ only. The both shifts in RLC (at 24°C and 42°C) were small. In IC, the notable shift was found at 24°C in barley and 42°C in maize (Fig. 9A,E). Under near lethal HS (46°C), LH conditions increased Y(I)/X(II) ratio in the both species under low and moderate AL intensities (Fig. 9G,H); IC analysis revealed a large increase of the ratio in maize (Fig. 9G).

Summarizing, LH conditions reduced amounts of PSI and PSII proportionally and decreased relative activity of PSII more than PSI (Fig. 9). The latter effect was small and found in control, genuine HS, and nearly lethal HS conditions at the relatively low intensities of light. The large shift in favor of PSI was found in maize under nearly lethal HS and light intensity to that plants were adapted during the experiment.

## Conclusion

The current research was designated for studying possible effect of air humidity on the photochemical activities of PSII and PSI under conditions of HS. The conditions of LH decreased activities of the both photosystems. The small effect was found in control conditions (24°C); plant growth was not influenced with this reduction. The moderate HS (37°C) decreased the maximal photochemical activities of PSII in the both species and PSI in barley; it was counterbalanced with the increase of relative photochemical activities of the both photosystems in the both species. The air RH conditions did influence nor activities of photosystems, neither plant growth. Under this temperature, the growth of barley was inhibited slightly; the growth of maize shoot was not retarded at all.

The temperature elevation to 42°C was followed with the decrease of the maximal photochemical activity of PSI in maize and some additional reduction of relative photochemical activities of PSII and PSI in barley. The growth of plants was retarded obviously. The conditions of LH reduced additionally both the activities of photosystems and plant growth; all the effects of air RH were small.

Under near lethal HS (46°C), the both activities of photosystems and plant growth were retarded greatly. The maximal activities of PSII and PSI were reduced much more. The up regulation of relative activities was exhausted; the relative photochemical activities of PSII and PSI were at the control level or below it. Some parameters of plant growth were reduced several times. The conditions of LH reduced additionally both the activities of photosystems and plant growth; all the effects of air RH were large. In LH conditions, barley plants were dying if the treatment was extended for 3-5 h. Maize plants increased water uptake by roots in LH conditions and survived.

Lower humidity of air decreased activities of the both photosystems. Probably, plants own mechanisms to alleviate this effect. Under severe HS, an additional impact of lower air humidity can be critical for plant survival.

## Supporting information

Supplementary Tables and Commentaries

## Supplementary data

File “Supplementary Tables and Commentaries”:

*Commentary 1*. Linear regression analysis of data presented in Fig. 6F and Supplementary Fig. S2 (42°C).

*Commentary 2*. Nonparametric analysis of data presented in Fig. 9A.

Supplementary Tables S1-S5 – complete set of data and statistics on the plant growth; the selected examples from this set are shown in Fig. 1.

*Table S1*. Root growth.

*Table S2*. Shoot growth.

*Table S3*. The second leave growth.

*Table S4*. The first leave growth.

*Table S5*. Stem growth.

*Table S6*. Water contents in plant organs. Complete set of data and statistics; reduced version is presented in Fig. 2.

*Table S7*. Water uptake by roots that is shown in Fig. 3. Complete statistics.

*Table S8*. Photosynthetic pigments in leaves. Complete set of data and statistics; reduced version is presented in Fig. 4.

*Table S9*. Basic photosynthetic values in the dark-adapted plants. Complete set of data and statistics; reduced version is presented in Fig. 5.

File “Supplementary Figures”:

*Fig. S1*. Fv/Fm dynamics. Revisualization of data from Fig. 5C.

*Fig. S2*. Dynamics of Φ_PSII_ in IC; visualization for tracing RH effect.

*Fig. S3*. Dynamics of photochemical coefficients of PSII in IC. Revisualization of the data from Fig. 6 and Supplementary Fig. S2 for tracing temperature effect.

*Fig. S4*. Dynamics of Φ_PSII_ in RLC; visualization for tracing RH effect.

*Fig. S5*. Dynamics of photochemical coefficients of PSII in RLC. Revisualization of the data from Fig. 7 and Supplementary Fig. S4 for tracing temperature effect.

*Fig. S6*. Dynamics of quantum yield of PSI. Revisualization of the data from Fig. 8 for tracing temperature effect.

*Fig. S7*. Dynamics of the ratio Y(I)/X(II). Revisualization of the data from Fig. 9 for tracing temperature effect.

## Acknowledgement

The authors thank Prof. A.V. Rubanovich (VIGG, RAS) for the additional statistic analysis (see, Supplementary Commentaries 1-2). The seeds of maize were kindly provided by Dr. E.V. Kartamysheva (VNIIMK, Don experimental station). The research was carried out within the state assignment of Ministry of Science and Higher Education of the Russian Federation (theme No.121040800153-1) and supported by the Russian Science Foundation grant (project No. 14-14-00584).

## Author contribution

EAL: conceptualization, methodology, project administration, some data acquisition with PAM, PAM data analysis, visualization, writing; MAK: most data acquisition with PAM; AAK: plant growing, all the data acquisition except PAM, preparation of tables, statistical analysis; NLP: data acquisition with PAM in a preliminary experiment critical for the design of model; VVK: funding acquisition, resources, supervision. All authors read and approved the manuscript. EAL is the author responsible for contact and ensures communication.

## Data availability

All data supporting the findings of this study are available within the paper and within its supplementary materials published online.

## Abbreviations

AL: actinic light
Chl: chlorophyll
HH: higher (relative) humidity (of air)
HS: heat stress
IC: induction curve
LH: lower (relative) humidity (of air)
PAM: pulse amplitude modulation
PSI, PSII: photosystem I and II
P_700_: reaction center Chl of PSI
RH: relative humidity (of air)
RLC: rapid light curve
SP: saturation pulse

## References

Alhaithloul HAS. 2019. Impact of Combined Heat and Drought Stress on the Potential Growth Responses of the Desert Grass Artemisia sieberi alba: Relation to Biochemical and Molecular Adaptation. Plants 8, 416. doi: 10.3390/plants8100416

Allakhverdiev SI, Kreslavski VD, Klimov VV, Los DA, Carpentier R, Mohanty P. 2008. Heat stress: an overview of molecular responses in photosynthesis. Photosynthesis Research 98, 541–550. doi: 10.1007/s11120-008-9331-0

Arena C, Conti S, Francesca S, Melchionna G, Hájek J, Barták M, Barone A, Rigano MM. 2020. Eco-Physiological Screening of Different Tomato Genotypes in Response to High Temperatures: A Combined Field-to-Laboratory Approach. Plants 9, 508. doi: 10.3390/plants9040508

Berry JA, Björkman O. 1980. Photosynthetic response and adaptation to temperature in higher plants. Annual Review of Plant Physiology 31, 491–543. doi: 10.1146/annurev.pp.31.060180.002423

Bukhov NG, Carpentier R. 2000. Heterogeneity of photosystem II reaction centers as influenced by heat treatment of barley leaves. Physiologia Plantarum 110, 279–285. doi: 10.1034/j.1399-3054.2000.110219.x

Camejo D, Rodrıguez P, Morales MA, Dell’Amico JM, Torrecillas A, Alarcon JJ. 2005. High temperature effects on photosynthetic activity of two tomato cultivars with different heat susceptibility. Journal of Plant Physiology 162, 281—289. doi: 10.1016/j.jplph.2004.07.014

Chávez-Arias CC, Ligarreto-Moreno GA, Ramírez-Godoy A, Restrepo-Díaz H. 2021. Maize Responses Challenged by Drought, Elevated Daytime Temperature and Arthropod Herbivory Stresses: A Physiological, Biochemical and Molecular View. Frontiers in Plant Science 12, article 702841. doi: 10.3389/fpls.2021.702841

Diaz M, de Haro V, Muňoz R, Quiles M. 2007. Chlororespiration is involved in the adaptation of Brassica plants to heat and high light intensity. Plant, Cell and Environment 30, 1578–1585. doi: 10.1111/j.1365-3040.2007.01735.x

Doğru A. 2021. Effects of heat stress on photosystem II activity and antioxidant enzymes in two maize cultivars. Planta 253, 85. doi: 10.1007/s00425-021-03611-6

Dual-PAM_1e. Instruction manual for DUAL-PAM-100. 2009. Heinz Walz GmbH. https://www.walz.com/files/downloads/manuals/dual-pam-100/Dual-PAM_1e.pdf new Accessed April 2022.

Feller U, Crafts-Brandner SJ, Salvucci ME. 1998. Moderately high temperatures inhibit ribulose-1,5-biphosphate carboxylase/oxygenase (Rubisco) activase-mediated activation of Rubisco. Plant Physiology 116, 539–546. doi:10.1104/pp.116.2.539

Ferguson JN, McAusland L, Smith KE, Price AH, Wilson ZA, Murchie EH. 2020. Rapid temperature responses of photosystem II efficiency forecast genotypic variation in rice vegetative heat tolerance. The Plant Journal 104, 839–855. doi: 10.1111/tpj.14956

Franco E, Alessandrelli S, Masojidek J, Margonelli A, Giardi MT. 1999. Modulation of D1 protein turnover under cadmium and heat stresses monitored by [35S]methionine incorporation. Plant Science 144, 53–61. doi: 10.1016/S0168-9452(99)00040-0

Genty B, Briantais J-M, Baker NR. 1989. The relationship between the quantum yield of photosynthetic electron transport and quenching of chlorophyll fluorescence. // Biochimica et Biophysica Acta 990, 87–92. doi: 10.1016/S0304-4165(89)80016-9

Goss R, Lohr M, Latowski D, Grzyb J, Vieler A, Wilhelm C, Strzalka K. 2005. Role of hexagonal structure-forming lipids in diadinoxanthin and violaxanthin solubilization and de-epoxidation. Biochemistry 44, 4028–4036. doi: 10.1021/bi047464k

Gounaris K, Brain ARR, Quinn PJ, Williams WP. 1984. Structural reorganization of chloroplast thylakoid membranes in response to heat stress. Biochimica et Biophysica Acta 766, 198–208. doi: 10.1016/0005-2728(84)90232-9

Havaux M. 1996. Short-term responses of Photosystem I to heat stress. Induction of a PS II-independent electron transport through PSI fed by stromal components. Photosynthesis Research 47, 85–97. doi: 10.1007/BF00017756

Havaux M, Greppin H, Strasser RJ. 1991. Functioning of photosystems I and II in pea leaves exposed to heat stress in the presence or absence of light. Analysis using in-vivo fluorescence, absorbance, oxygen and photoacoustic measurements. Planta 186, 88–98. doi: 10.1007/BF00201502

Havaux M, Strasser RJ. 1990. Protection of Photosystem II by Light in Heat-Stressed Pea Leaves. Zeitschrift für Naturforschung 45c, 1133–1141. https://zfn.mpdl.mpg.de/data/Reihe_C/45/ZNC-1990-45c-1133.pdf

Horvath I, Glatz A, Varvasovszki V, et al. (1998) Membrane physical state controls the signaling mechanism of the heat shock response in Synechocystis PCC 6803: identification of hsp17 as a fluidity gene. Proceedings of the National Academy of Sciences of USA 95, 3513–3518. doi:10.1073/pnas.95.7.3513

Kalaji HM, Schansker G, Ladle RJ, et al. 2014. Frequently asked questions about in vivo chlorophyll fluorescence: practical issues. Photosynthesis Research 122, 121–158. doi: 10.1007/s11120-014-0024-6

Kalituho LN, Pshybytko NL, Kabashnikova LF, Jahns P. 2003. Photosynthetic apparatus and high temperature: Role of light. Bulgarian Journal of Plant Physiology Special Issue, 281–289. http://www.bio21.bas.bg/ipp/gapbfiles/essa-03/03_essa_281-289.pdf

Klughammer C, Schreiber U. 1994. An improved method, using saturating light pulses, for the determination of photosystem I quantum yield via P700+-absorbance changes at 830 nm. Planta 192, 261–268. doi: 10.1007/BF01089043

Killi D, Bussotti F, Raschi A, Haworth M. 2017. Adaptation to high temperature mitigates the impact of water deficit during combined heat and drought stress in C3 sunflower and C4 maize varieties with contrasting drought tolerance. Physiologia Plantarum 159, 130–147. doi: 10.1111/ppl.12490

Killi D, Raschi A, Bussotti F. 2020. Lipid Peroxidation and Chlorophyll Fluorescence of Photosystem II Performance during Drought and Heat Stress is Associated with the Antioxidant Capacities of C3 Sunflower and C4 Maize Varieties. International Journal of Molecular Sciences 21, 4846. doi: 10.3390/ijms21144846

Lichtenthaler HK. 1987. Chlorophylls and carotenoids: pigments of photosynthetic biomembranes. Methods in Enzymology 148, 350–382. doi: 10.1016/0076-6879(87)48036-1

Lichtenthaler HK, Buschmann C, Knapp M. 2005. How to correctly determine the different chlorophyll fluorescence parameters and the chlorophyll fluorescence decrease ratio RFd of leaves with the PAM fluorometer. Photosynthetica 43, 379–393. doi: 10.1007/s11099-005-0062-6

Lobell DB, Asner GP. 2003. Climate and management contributions to recent trends in U.S. agricultural yields. Science 299, 1032. doi: 10.1126/science.1078475

Lysenko EA. 2021. Application of fast light-readapted plants for measurement of chlorophyll fluorescence and P700 light absorption with the RLC method. Photosynthetica 59, 245–255. doi: 10.32615/ps.2021.015

Lysenko EA, Klaus AA, Kartashov AV, Kusnetsov VV. 2019. Distribution of Cd and other cations between the stroma and thylakoids: a quantitative approach to the search for Cd targets in chloroplasts. Photosynthesis Research 139, 337–358. doi: 10.1007/s11120-018-0528-6

Lysenko EA, Klaus AA, Kartashov AV, Kusnetsov VV. 2020. Specificity of Cd, Cu, and Fe effects on barley growth, metal contents in leaves and chloroplasts, and activities of photosystem I and photosystem II. Plant Physiology and Biochemistry 147, 191–204. doi: 10.1016/j.plaphy.2019.12.006

Monneveux P, Pastenes C, Reynolds MP. (2003) Limitations to photosynthesis under light and heat stress in three high-yielding wheat genotypes. // J. Plant Physiol. 160. 657–666. doi: 10.1078/0176-1617-00772

Murakami Y, Tsuyama M, Kobayashi Y, Kodama H, Iba K. 2000. Trienoic fatty acids and plant tolerance of high temperature. Science 287, 476–479. doi: 10.1126/science.287.5452.476

Nellaepalli S, Mekala NR, Zsiros O, Mohanty P, Subramanyam R. 2011. Moderate heat stress induces state transitions in Arabidopsis thaliana. Biochimica et Biophysica Acta 1807, 1177–1184. doi: 10.1016/j.bbabio.2011.05.016

Pastenes C, Horton R. 1996. Effect of high temperature on photosynthesis in beans. Plant Physiology 112, 1245–1251. doi: 10.1104/pp.112.3.1253

Pshybytko NL, Kruk J, Kabashnikova LF, Strzalka K. 2008. Function of plastoquinone in heat stress reactions of plants. Biochimica et Biophysica Acta 1777, 1393–1399. doi:10.1016/j.bbabio.2008.08.005

Radin JW, Lu Z, Percy RG, Zeiger E. 1994. Genetic variability for stomatal conductance in Pima cotton and its relation to improvements of heat adaptation. Proceedings of the National Academy of Sciences of USA 91, 7217–7221. doi: 10.1073/pnas.91.15.7217

Sato N, Sonoike K, Kawaguchi A, Tsuzuki M. 1996. Contribution of lowered unsaturation levels of chloroplast lipids to high temperature tolerance of photosynthesis in Chlamydomonas reinhardtii. Journal of Photochemistry and Photobiology B: Biology 36, 333–337. doi: 10.1016/S1011-1344(96)07389-7

Schreiber U, Schliwa U, Bilger W. 1986. Continuous recording of photochemical and nonphotochemical chlorophyll fluorescence quenching with a new type of modulation fluorometer. Photosynthesis Research 10, 51–62. doi: 10.1007/BF00024185

Sharkey TD. 2005. Effects of moderate heat stress on photosynthesis: importance of thylakoid reactions, rubisco deactivation, reactive oxygen species, and thermotolerance provided by isoprene. Plant, Cell and Environment 28, 269–277. doi: 10.1111/j.1365-3040.2005.01324.x

Sharma DK, Andersen SB, Ottosen C-O, Rosenqvist E. 2015. Wheat cultivars selected for high Fv/Fm under heat stress maintain high photosynthesis, total chlorophyll, stomatal conductance, transpiration and dry matter. Physiologia Plantarum 153, 284–298. doi: 10.1111/ppl.12245

Speck O, Schlechtendahl M, Borm F, Kampowski T, Speck T. 2018. Humidity-dependent wound sealing in succulent leaves of Delosperma cooperi – An adaptation to seasonal drought stress. Beilstein Journal of Nanotechnology 9, 175–186. doi:10.3762/bjnano.9.20

Tanaka Y, Nishiyama Y, Murata N. 2000. Acclimation of the Photosynthetic Machinery to High Temperature in Chlamydomonas reinhardtii Requires Synthesis de Novo of Proteins Encoded by the Nuclear and Chloroplast Genomes. Plant Physiology 124, 441–449. doi: 10.1104/pp.124.1.441

Turan S, Kask K, Kanagendran A, Li S, Anni R, Talts E, Rasulov B, Kännaste A, Niinemets Ü. 2019. Lethal heat stress-dependent volatile emissions from tobacco leaves: what happens beyond the thermal edge? Journal of Experimental Botany 70, 5017–5030. doi:10.1093/jxb/erz255

van Kooten O, Snel JFH. 1990. The use of chlorophyll fluorescence nomenclature in plant stress physiology. Photosynthesis Research 25, 147–150. doi: 10.1007/BF00033156

Wahid A, Gelani S, Ashraf M, Foolad MR. 2007. Heat tolerance in plants: an overview. Environmental and Experimental Botany 61, 199–223. doi: 10.1016/j.envexpbot.2007.05.011

Wang L, Ma K-B, Lu Z-G, Ren S-X, Jiang H-R, Cui J-W, Chen G, Teng N-J, Lam H-M, Jin B. 2020. Differential physiological, transcriptomic and metabolomic responses of Arabidopsis leaves under prolonged warming and heat shock. BMC Plant Biology 20, 86. doi: 10.1186/s12870-020-2292-y

Wise RR, Olson AJ, Schrader SM, Sharkey TD. 2004. Electron transport is the functional limitation of photosynthesis in field-grown Pima cotton plants at high temperature. Plant, Cell and Environment 27, 717–724. doi: 10.1111/j.1365-3040.2004.01171.x

Yamane Y, Shikanai T, Kashino Y, Koike H, Satoh K. 2000. Reduction of Q_A_ in the dark: Another cause of fluorescence Fo increases by high temperatures in higher plants. Photosynthesis Research 63, 23–34. doi: 10.1023/A:1006350706802

Yan K, Chen P, Shao H, Shao C, Zhao S, Brestic M. 2013. Dissection of Photosynthetic Electron Transport Process in Sweet Sorghum under Heat Stress. PLoS ONE 8, e62100. doi:10.1371/journal.pone.0062100

Yudina L, Sukhova E, Gromova E, Nerush V, Vodeneev V, Sukhov V. 2020. A light-induced decrease in the photochemical reflectance index (PRI) can be used to estimate the energy-dependent component of non-photochemical quenching under heat stress and soil drought in pea, wheat, and pumpkin. Photosynthesis Research, 146, 175–187. doi: 10.1007/s11120-020-00718-x

Zhou R, Yu X, Li X, dos Santos TM, Rosenqvist E, Ottosen C-O. 2020. Combined high light and heat stress induced complex response in tomato with better leaf cooling after heat priming. Plant Physiology and Biochemistry 151, 1–9. doi: 10.1016/j.plaphy.2020.03.0111

Zubo YaO, Lysenko EA, Aleinikova AYu, Kusnetsov VV, Pshibytko NL. 2008. Changes in the Transcriptional Activity of Barley Plastome Genes under Heat Shock. Russian Journal of Plant Physiology 55, 293–300. doi: 10.1134/S1021443708030011

