## Supplementary Tables and Commentaries for "Lower air humidity reduced both the plant growth and activities of photosystems I and II under prolonged heat stress"

**Source:** bioRxiv

### Supplementary Commentaries

**Commentary 1.**

Under the temperature 42°C, maize (Zm) plants demonstrated higher level of PSII activities at HH conditions than LH conditions. According to the coefficient qP, the difference was negligible (Fig. 6E); it was not analyzed further. According to the coefficient X(II) and $\Phi_{PSII}$, the difference was notable; however, the pairwise comparisons of the individual data points with the use of t-test failed to reveal significant difference for the both X(II) (Fig. 6E) and $\Phi_{PSII}$ (Suppl. Fig. S2 42°C). Therefore, the whole dynamics were compared with the use of z-test.

The measurements in dark adapted state and 1 s under the conditions of AL demonstrated very high and low values correspondingly. After 41 s under the conditions of AL, dynamics of the both coefficients reached steady state levels with a slow further growth. The range from 41 s to 441 s under the AL comprised 11 measurement points. The ranges 41-441 s were used for calculating of equations of linear regression; the equations were compared with the use of z-test.

**X(II) (Fig. 6F)**

Equations of linear regression:

Zm HH: $y=(0.646 +/- 0.006) + (2.8e^{-4} +/- 2.1e^{-5})x$

Zm HL: $y=(0.566 +/- 0.006) + (2.5e^{-4} +/- 2.3e^{-5})x$

The slopes do not differ significantly. For the free members of equations:

$Z = (0.646-0.566) / \sqrt{}((0.006)^2+(0.006)^2) = 9.43$, where √ denotes “root square”.

**The difference is significant at $p < 1e^{-13}$.**

**$\Phi_{PSII}$ (Suppl. Fig. S2 42°C)**

Equations of linear regression:

Zm HH: $y=(0.596 +/- 0.008) + (1.9e^{-4} +/- 3.1e^{-5})x$

Zm HL: $y=(0.555 +/- 0.006) + (2.0e^{-4} +/- 2.2e^{-5})x$

The slopes do not differ significantly. For the free members of equations:

$Z = (0.596-0.555) / \sqrt{} ((0.008)^2+(0.006)^2) = 4.1$, where √ denotes “root square”.

**The difference is significant at $p = 2.1e^{-5}$ (<0,015).**

The linear regression analysis was kindly provided by Prof. A.V. Rubanovich (Vavilov Institute of General Genetics, RAS)

**Commentary 2.**

In control, the ratio Y(I)/X(II) was significantly different in HH and LH variants of barley plants. In maize, the difference was smaller and insignificant according to pairwise comparison of the individual data points with the use of t-test (Fig. 9A). However, the steady state level of the ratio Y(I)/X(II) (41-441 s at AL) was always higher in ZmLH than ZmHH variants. The range from 41 s to 441 s under the AL comprised 11 measurement points. The external points at 41 s and 441 s are nearly equal and; they were rejected. The rest 9 points (81-401 s of AL) demonstrated clearly ZmLH > ZmHH (Fig. 9A).

The probability that ZmLH > ZmHH in each point is ½. The probability that ZmLH > ZmHH in all the 9 point is equal to $(½)^9 = 0{,}001953$; this is one-tail paired nonparametric binomial test. In the two-tailed test, the probability is 0.003906.

Under the control temperature (24°C), the steady state level of **the ratio Y(I)/X(II) is significantly higher in ZmLH than ZmHH (Fig. 9A) at p<0.004**. The difference is significant according to the two-tailed paired nonparametric binomial test; the difference between the absolute values of means is very small.

The author thanks Prof. A.V. Rubanovich for the advice to apply this nonparametric test.

#### Supplementary Tables

**Table S1.** Length and weight of roots before the experiment (7 day) and after growth under control or elevated temperatures at contrast conditions of air relative humidity (9 day).

| Roots | Temp. °C | Plant age 7-day | Plant age 9-day HH | Plant age 9-day LH | Increment HH | Increment LH |
|---|---|---|---|---|---|---|
| | | | HH | LH | HH | LH |
| *Zea mays* | | | | | | |
| Length, cm | 24 | 17.7±0.7 a 1 | 20.5±0.6 b 12 | 20.9±0.7 b 1 | 2.74±0.29 1 | 3.23±1.12 1 |
| | 37 | 17.7±0.8 a 1 | 21.8±0.6 b 1 | 20.7±0.6 b 1 | 4.08±1.07 1 | 3.04±1.49 1 |
| | 42 | 17.9±0.8 a 1 | 18.9±0.6 ab 23 | 19.9±0.6 b 12 | 1.03±0.76 12 | 2.03±0.89 1 |
| | 46 | 18.0±0.9 a 1 | 17.9±0.7 a 3 | 18.5±0.9 a 2 | -0.04±0.80 2 | 0.54±1.36 1 |
| DW, mg | 24 | 7.8 ±0.6 a 1 | 9.6±0.6 a 1 | 10.2±0.9 a 1 | 1.8±0.8 1 | 2.5±0.8 1 |
| | 37 | 6.6 ± 0.7 a 1 | 8.4±0.5 a 1 | 7.4±0.2 a 2 | 1.8±0.9 1 | 0.9±0.9 1 |
| | 42 | 6.8 ± 0.6 a 1 | 7.9±0.5 a 1 | 7.8±0.4 a 2 | 1.2±0.6 1 | 1.0±0.6 1 |
| | 46 | 7.7 ± 0.6 a 1 | 5.8±0.4 c 2 | 5.2±0.3 c 3 | -1.9±0.5 2 | -2.5±0.8 2 |
| FW, mg | 24 | 135 ±10 a 1 | 148±11 ab 1 | 165±10 b 1 | 13.0±19.1 1 | 29.8±16.8 1 |
| | 37 | 111±7 a 2 | 133±7 b 1 | 127±9 ab 2 | 21.9±15.8 1 | 16.3±13.7 1 |
| | 42 | 111±7 a 2 | 112±5 a 2 | 112±8 a 2 | 0.7±11.8 1 | 0.4±9.8 1 |
| | 46 | 125±8 a 12 | 79±5 c 3 | 71±6 c 3 | -45.2±8.8 2 | -53.9±11.7 2 |
| *Hordeum vulgare* | | | | | | |
| Length, cm | 24 | 7.1±0.6 a 3 | 10.0±0.5 b 12 | 9.7±0.4 b 1 | 2.89±0.20 1 | 2.59±0.79 1 |
| | 37 | **9.9**±0.4 a 1 | 10.0±0.3 a 1 | 9.7±0.3 a 1 | 0.13±0.30 2 | -0.19±0.47 2 |
| | 42 | 7.5±0.4 a 23 | 9.1±0.4 b 2 | 8.8±0.4 b 12 | 1.52±0.60 12 | 1.20±0.74 12 |
| | 46 | 8.3±0.4 ab 2 | 9.3±0.5 b 12 | 8.1±0.3 a 2 | 1.00±0.32 2 | 0.05±0.59 2 |
| DW, mg | 24 | 5.5±0.4 a 1 | 6.5±0.4 a 12 | 6.3±0.1 a 1 | 0.9±0.7 12 | 0.8±0.5 1 |
| | 37 | 5.9±0.1 a 1 | 6.9±0.2 b 1 | 6.8±0.4 ab 1 | 1.0±0.3 1 | 0.8±0.5 1 |
| | 42 | 6.0±0.1 a 1 | 5.4±0.2 a 2 | 5.6±0.2 a 2 | -0.5±0.1 2 | -0.3±0.2 1 |
| | 46 | 5.8±0.2 a 1 | 4.4±0.2 c 3 | 4.1±0.2 c 3 | -1.4±0.2 3 | -1.7±0.3 2 |
| FW, mg | 24 | 60±3 a 2 | 75±4 b 1 | 74±3 b 1 | 15.1±5.6 1 | 13.9±3.7 1 |
| | 37 | 70±3 a 1 | 70±3 a 1 | 70±4 a 1 | 0.5±5.4 12 | 0.1±4.9 12 |
| | 42 | 64±4 a 12 | 50±2 c 2 | 54±3 ac 2 | -14.0±3.3 23 | -10.0±4.5 2 |
| | 46 | 67±3 a 12 | 45±3 c 2 | 37±1 d 3 | -21.3±3.1 3 | -29.1±4.2 3 |

Pots with 7-day old plants were placed to thermostat chambers for 48 h. Just before the placement, a number of plants were taken from each pot for the measurement of sizes, masses, and contents of water and photopigments. After 48 h, another portion of plants were taken from the same pots and the same parameters were measured. For parameters of grows (sizes & masses), the difference between 7-day and 9-day old plants was considered as "Increment". Before the experiment, all plants were grown in same conditions and are equal. The difference between 7-day old plants' mean values was happened by chance bias in a sample formation; it was statistically significant in some cases. In such cases, it is better to consider all four means of 7-day old plants and find a general level. The most different values are marked with red font.
"Root length" is the distance between a caryopses and a most distant point of roots.
HH – higher (relative) humidity of air, LH - lower (relative) humidity of air, DW – dry weight, FW – fresh weight. Data are presented as means ± standard error (SE). n.d. – not determined.
1-4 – differences between variants of temperature regime are significant at $p \leq 0.05$.
a-e – differences between plants of a same temperature regime before (7d) and after experiment (9d) at different conditions of air relative humidity (HH & LH) are significant at $p \leq 0.05$.
* - difference between increments at HH and LH conditions (at a same temperature) is significant at $p \leq 0.05$.

**Table S2.** Height and weight of shoots before the experiment (7 day) and after growth under control or elevated temperatures at contrast conditions of air relative humidity (9 day).

| Shoot | Temp. °C | Plant age 7-day | Plant age 9-day HH | Plant age 9-day LH | Increment HH | Increment LH |
|---|---|---|---|---|---|---|
| *Zea mays* | | | | | | |
| Height, cm | 24 | 16.0 ± 0.7 a 12 | 21.7 ± 0.7 b 1 | 21.3 ± 0.5 b 1 | 5.6±1.3 1 | 5.2±1.0 1 |
| | 37 | 15.6 ± 0.6 a 2 | 22.5 ± 0.7 b 1 | 22.0 ± 0.6 b 1 | 6.9±1.1 1 | 6.4±0.8 1 |
| | 42 | 16.7 ± 0.5 a 12 | 20.6 ± 0.6 b 1 | 21.1 ± 0.6 b 1 | 4.0±1.2 12 | 4.5±0.5 1 |
| | 46 | 17.7 ± 0.6 a 1 | 18.7 ± 0.5 a 2 | 18.5 ± 0.6 a 2 | 1.0±0.6 2 | 0.8±0.8 2 |
| DW, mg | 24 | 28±2 a 1 | 43±4 b 12 | 40±2 b 1 | 15±5 1 | 14±3 1 |
| | 37 | 26±3 a 1 | 45±2 b 1 | 41±1 b 1 | 18±5 1 | 14±3 1 |
| | 42 | 28±1 a 1 | 41±4 b 12 | 41±3 b 1 | 13±3 1 | 13±3 1 |
| | 46 | 29±1 a 1 | 38±2 b 2 | 38±2 b 1 | 8±1 1 | 9±3 1 |
| FW, mg | 24 | 383 ± 22 a 1 | 549 ± 40 b 1 | 528 ± 24 b 1 | 167 ± 69 1 | 145 ± 39 1 |
| | 37 | 362 ± 22 a 1 | 507 ± 23 b 1 | 477 ± 24 b 12 | 144 ± 58 1 | 114 ± 44 1 |
| | 42 | 386 ± 14 a 1 | 426 ± 23 a 2 | 427 ± 22 a 2 | 41 ± 39 12 | 41 ± 25 1 |
| | 46 | 400 ± 15 a 1 | 385 ± 17 a 2 | 326 ± 14 c 3 | -15 ± 11 2* | -74 ± 21 2* |
| *Hordeum vulgare* | | | | | | |
| Height, cm | 24 | 15.6±0.4 a 12 | 18.6±0.6 b 1 | 17.2±0.4 b 1 | 2.99±0.17 1 | 1.66±0.51 1 |
| | 37 | 15.6±0.2 a 12 | 16.5±0.2 b 12 | 16.8±0.2 b 1 | 0.94±0.29 2 | 1.18±0.41 1 |
| | 42 | 15.3±0.3 a 2 | 16.0±0.2 b 3 | 16.0±0.2 b 2 | 0.68±0.29 2 | 0.67±0.48 1 |
| | 46 | 16.1±0.3 a 1 | 16.5±0.2 a 2 | n.d. | 0.38±0.20 2 | n.d. |
| DW, mg | 24 | 21±2 a 1 | 31±1 b 12 | 29±0 b 2 | 10±2 12 | 9±2 12 |
| | 37 | 20±0 a 1 | 30±1 b 2 | 31±2 b 12 | 10±1 2 | 11±2 12 |
| | 42 | 20±1 a 1 | 31±1 b 12 | 33±1 b 1 | 13±0 1 | 12±1 1 |
| | 46 | 21±1 a 1 | 33±1 e 1 | 28±1 b 2 | 12±1 12* | 7±2 2* |
| FW, mg | 24 | 232±9 a 1 | 315±13 b 1 | 310±10 b 1 | 83±19 1 | 78±16 1 |
| | 37 | 245±7 a 1 | 284±7 b 2 | 287±10 b 1 | 39±7 1 | 42±13 12 |
| | 42 | 230±10 a 1 | 269±8 b 2 | 253±7 ab 2 | 38±5 1 | 21±10 2 |
| | 46 | 244±10 a 1 | 170±10 c 3 | 61±3 d 2 | -73±18 2* | -180±18 3* |

Plant height is the distance between a caryopses and a highest point of shoot.
All other designations are the same as in Table S1.

**Table S3.** Size and weight of the 2nd leaves before the experiment (7 day) and after growth under control or elevated temperatures at contrast conditions of air relative humidity (9 day).

| 2nd leaf | Temp. °C | Plant age 7-day | 9-day HH | 9-day LH | Increment HH | Increment LH |
|---|---|---|---|---|---|---|
| *Zea mays* | | | | | | |
| Length, cm | 24 | 10.5 ± 0.6 a 12 | 15.3 ± 0.5 b 1 | 14.9 ±0.4 b 1 | 4.8±1.2 1 | 4.4±1.1 1 |
| | 37 | 9.7 ± 0.4 a 2 | 15.7 ± 0.6 b 1 | 15.1 ±0.5 b 1 | 6.0±0.5 1 | 5.5±0.4 1 |
| | 42 | 10.9 ± 0.5 a 12 | 14.4 ± 0.7 b 12 | 14.8 ±0.6 b 1 | 3.4±1.2 12 | 3.9±0.8 1 |
| | 46 | 11.9 ± 0.5 a 1 | 12.9 ± 0.4 a 2 | 13.0 ±0.5 a 2 | 1.0±0.5 2 | 1.1±0.8 2 |
| Width, cm | 24 | 1.15±0.03 a 1 | 1.13±0.03 a 1 | 1.14±0.03 a 1 | -0.02±0.05 1 | -0.01±0.04 1 |
| | 37 | 1.10±0.03 a 1 | 1.08±0.03 a 12 | 1.08±0.03 a 1 | -0.02±0.04 1 | -0.01±0.05 1 |
| | 42 | 1.13±0.03 a 1 | 1.03±0.03 c 2 | 1.11±0.03 a 1 | -0.10±0.06 1 | -0.02±0.06 1 |
| | 46 | 1.08±0.03 a 1 | 1.05±0.03 a 2 | 0.96±0.05 c 2 | -0.03±0.03 1 | -0.12±0.06 1 |
| DW, mg | 24 | 10.2±1.4 a 1 | 16.5±1.1 b 12 | 15.3±0.3 b 2 | 6.3±2.1 12 | 5.1±1.7 1 |
| | 37 | 8.7±0.8 a 1 | 17.8±0.4 e 1 | 16.4±0.4 b 1 | 9.1±1.2 1 | 7.7±0.8 1 |
| | 42 | 10.5±0.6 a 1 | 15.8±1.5 b 12 | 16.9±1.6 b 12 | 5.3±1.6 12 | 6.4±1.4 1 |
| | 46 | 10.9±0.7 a 1 | 15.5±0.8 b 2 | 17.0±1.1 b 12 | 4.5±0.4 2 | 6.1±1.1 1 |
| FW, mg | 24 | 119±9 a 12 | 175±10 b 1 | 170±7 b 1 | 56±22 1 | 52±20 1 |
| | 37 | 101±7 a 2 | 168±7 b 1 | 161±7 b 1 | 67±12 1 | 60±11 1 |
| | 42 | 120±6 a 12 | 142±9 b 2 | 154±10 b 1 | 22±19 12 | 34±18 1 |
| | 46 | 127±6 a 1 | 121±6 a 3 | 97±6 c 2 | -6±5 2 | -29±9 2 |
| *Hordeum vulgare* | | | | | | |
| Length, cm | 24 | 5.0±0.3 a 1 | 12.9±0.6 b 1 | 11.5±0.4 b 1 | 7.9±0.5 1 | 6.5±0.4 1 |
| | 37 | 4.7±0.2 a 1 | 9.4±0.4 b 2 | 9.5±0.2 b 2 | 4.7±0.4 2 | 4.8±0.3 2 |
| | 42 | 4.7±0.2 a 1 | 7.8±0.3 b 3 | 7.4±0.3 b 3 | 3.1±0.4 3 | 2.7±0.3 3 |
| | 46 | 5.2±0.3 a 1 | 6.2±0.2 b 4 | n.d. | 1.0±0.3 4 | n.d. |
| Width, cm | 24 | 0.39±0.02 a 1 | 0.63±0.02 b 1 | 0.63±0.01 b 1 | 0.24±0.03 1 | 0.24±0.02 1 |
| | 37 | 0.38±0.01 a 1 | 0.49±0.02 b 2 | 0.49±0.02 b 2 | 0.10±0.02 2 | 0.11±0.01 2 |
| | 42 | 0.39±0.01 a 1 | 0.51±0.02 b 2 | 0.48±0.02 b 2 | 0.13±0.03 2 | 0.10±0.02 2 |
| | 46 | 0.39±0.02 a 1 | 0.41±0.02 a 3 | n.d. | 0.01±0.02 3 | n.d. |
| DW, mg | 24 | 2.0±0.5 a 1 | 9.8±0.8 b 1 | 8.7±0.2 b 1 | 7.8±0.7 1 | 6.6±0.4 1 |
| | 37 | 1.8±0.1 a 1 | 6.6±0.4 b 2 | 6.7±0.3 b 2 | 4.9±0.5 2 | 5.1±0.3 2 |
| | 42 | 1.8±0.2 a 1 | 5.5±0.7 b 2 | 5.1±0.7 b 2 | 4.3±0.1 2 | 3.5±0.9 2 |
| | 46 | 2.1±0.2 a 1 | 5.3±0.5 b 2 | n.d. | 3.2±0.5 2 | n.d. |
| FW, mg | 24 | 19±2 a 1 | 79±7 b 1 | 77±4 b 1 | 61±8 1 | 58±4 1 |
| | 37 | 19±2 a 1 | 52±4 b 2 | 51±3 b 2 | 33±3 2 | 33±1 2 |
| | 42 | 18±2 a 1 | 43±3 b 2 | 36±3 b 3 | 24±4 2 | 18±3 3 |
| | 46 | 20±2 a 1 | 19±2 a 3 | n.d. | -1±2 3 | n.d. |

"2nd leaf" — leaf blade only.
All designations are the same as in Tables S1.

**Table S4.** Size and weight of the 1st leaves before the experiment (7 day) and after growth under control or elevated temperatures at contrast conditions of air relative humidity (9 day).

| 1st leaf | Temp. °C | Plant age 7-day | 9-day HH | 9-day LH | Increment HH | Increment LH |
|---|---|---|---|---|---|---|
| *Zea mays* | | | | | | |
| Length, cm | 24 | 5.1±0.2 a 1 | 5.7±0.2 2 b 1 | 5.3±0.2 a 12 | 0.6±0.2 1 | 0.3±0.3 1 |
| | 37 | 5.9±0.4 a 1 | 5.3±0.2 12 a 12 | 4.9±0.2 c 2 | -0.6±0.6 12 | -1.0±0.4 2 |
| | 42 | 5.4±0.2 a 1 | 5.3±0.3 12 a 12 | 5.8±0.3 a 1 | -0.2±0.2 2 | 0.3±0.5 12 |
| | 46 | 4.9±0.3 a 1 | 5.1±0.2 1 a 2 | 4.9±0.2 a 2 | 0.2±0.3 12 | -0.1±0.3 12 |
| Width, cm | 24 | 1.37±0.03 a 1 | 1.39±0.03 a 1 | 1.40±0.04 a 1 | 0.02±0.06 1 | 0.03±0.08 1 |
| | 37 | 1.35±0.03 a 1 | 1.38±0.03 a 1 | 1.34±0.04 a 12 | 0.03±0.03 1 | -0.01±0.01 1 |
| | 42 | 1.32±0.03 a 1 | 1.25±0.04 a# 2 | 1.32±0.04 a 12 | -0.07±0.05 1 | -0.01±0.03 1 |
| | 46 | 1.30±0.03 a 1 | 1.29±0.03 a 2 | 1.22±0.04 a# 2 | 0.00±0.05 1 | -0.08±0.06 1 |
| | Aver. | 1.34±0.01 # | | | | |
| DW, mg | 24 | 6.7±0.2 a 1 | 7.5±0.3 b 1 | 7.6±0.5 b 12 | 0.8±0.4 1 | 0.8±0.8 12 |
| | 37 | 7.8±1.0 ab 1 | 7.3±0.2 b 1 | 6.4±0.3 a 2 | -0.4±1.0 1 | -1.4±0.7 2 |
| | 42 | 7.0±0.4 a 1 | 7.0±0.2 a 1 | 7.8±0.4 a 1 | 0.0±0.5 1 | 0.9±0.1 1 |
| | 46 | 6.5±0.4 a 1 | 7.2±0.5 ab 1 | 7.8±0.2 b 1 | 0.7±0.6 1 | 1.4±0.5 1 |
| FW, mg | 24 | 77±4 a 1 | 88±4 a 1 | 86±4 a 1 | 11±8 1 | 10±9 1 |
| | 37 | 89±8 a 1 | 82±4 a 12 | 72±3 a 2 | -6±11 12 | -17±8 12 |
| | 42 | 84±4 a 1 | 72±5 a 23 | 81±4 a 12 | -12±5 2 | -3±5 12 |
| | 46 | 74±4 a 1 | 71±4 a 3 | 59±3 c 3 | -3±5 12 | -14±5 2 |
| *Hordeum vulgare* | | | | | | |
| Length, cm | 24 | 10.5±0.18 a 1 | 10.2±0.21 a 2 | 10.4±0.12 a 2 | -0.2±0.38 1 | -0.1±0.20 1 |
| | 37 | 10.7±0.15 a 1 | 10.7±0.16 a 12 | 10.9±0.18 a 1 | 0.1±0.29 1 | 0.2±0.29 1 |
| | 42 | 10.5±0.17 a 1 | 10.4±0.13 a 12 | 10.6±0.18 a 12 | -0.0±0.27 1 | 0.1±0.41 1 |
| | 46 | 10.8±0.12 a 1 | 10.8±0.14 a 1 | n.d. | -0.0±0.06 1 | n.d. |
| Width, cm | 24 | 0.78±0.02 a 1 | 0.76±0.01 a 2 | 0.75±0.01 a 2 | -0.02±0.01 1 | -0.03±0.02 1 |
| | 37 | 0.79±0.01 a 1 | 0.80±0.01 a 1 | 0.81±0.01 a 1 | 0.01±0.01 1 | 0.01±0.01 1 |
| | 42 | 0.78±0.01 a 1 | 0.78±0.02 a 12 | 0.79±0.01 a 1 | 0.00±0.01 1 | 0.01±0.02 1 |
| | 46 | 0.79±0.01 a 1 | 0.69±0.02 c 3 | n.d. | -0.09±0.03 2 | n.d. |
| DW, mg | 24 | 11.9±0.5 a 1 | 12.0±0.6 a 1 | 12.0±0.4 a 2 | 0.2±1.0 1 | 0.1±0.8 2 |
| | 37 | 12.4±0.1 a 1 | 13.8±0.2 b 2 | 14.2±0.9 ab 23 | 1.3±0.0 1 | 2.3±1.2 23 |
| | 42 | 11.7±0.3 a 1 | 16.6±0.4 b 3 | 17.1±1.0 b 3 | 5.2±0.5 2 | 5.4±1.2 3 |
| | 46 | 12.1±0.5 a 1 | 19.7±0.6 b 4 | 3.7±0.5 c 1 | 7.6±0.6 3* | -8.3±0.5 1* |
| FW, mg | 24 | 125±4 a 1 | 122±6 a 12 | 124±4 a 2 | -3±9 1 | -1±9 1 |
| | 37 | 135±5 a 1 | 135±4 a 1 | 137±5 a 1 | 0±3 1 | 2±7 1 |
| | 42 | 124±5 a 1 | 119±4 a 2 | 119±5 a 2 | -5±5 1 | -6±9 1 |
| | 46 | 129±5 a 1 | 71±6 c 3 | 25±2 d 3 | -58±10 2* | -105±4 2* |

"1st leaf" — leaf blade only.
Aver. – average mean ± SE calculated from all 7-day old plants (pots destined for further treatment at 24-46°C).
### - the difference from the average value of all 7-day-old plants is significant ($p<0.05$). See legend at the Table S1.
All other designations are the same as in Tables S1.

**Table S5.** Weight of stem with leaf sheaths before the experiment (7 day) and after growth under control or elevated temperatures at contrast conditions of air relative humidity (9 day).

| Stem | Temp. °C | Plant age 7-day | 9-day HH | 9-day LH | Increment HH | Increment LH |
|---|---|---|---|---|---|---|
| *Zea mays* | | | | | | |
| DW, mg | 24 | 10.9±0.9 a 1 | 18.7±2.7 b 12 | 17.0±1.8 b 12 | 7.8±3.1 12 | 6.6±1.2 12 |
| | 37 | 9.8±1.0 a 1 | 19.4±1.6 b 1 | 17.8±0.6 b 1 | 9.6±2.6 1 | 8.0±1.5 1 |
| | 42 | 10.5±0.5 a 1 | 18.3±2.0 b 12 | 16.2±2.0 b 12 | 7.7±1.7 1 | 5.6±1.6 12 |
| | 46 | 11.9±0.7 a 1 | 14.8±0.9 b 2 | 13.2±1.5 ab 2 | 3.0±0.3 2 | 1.4±2.0 2 |
| FW, mg | 24 | 187±11 a 1 | 287±28 b 1 | 271±20 b 1 | 100±43 1 | 84±17 1 |
| | 37 | 173±11 a 1 | 256±18 b 12 | 243±17 b 1 | 84±35 1 | 71±26 12 |
| | 42 | 182±7 a 1 | 213±15 a 23 | 192±14 a 2 | 31±17 12 | 10±10 23 |
| | 46 | 200±11 a 1 | 194±11 a 3 | 169±13 a 2 | -6±9 2 | -31±17 3 |
| *Hordeum vulgare* | | | | | | |
| DW, mg | 24 | 6.8±0.7 a 1 | 9.0±0.3 b 1 | 8.7±0.3 b 1 | 2.2±0.9 1 | 1.8±0.5 1 |
| | 37 | 6.4±0.2 a 1 | 9.7±0.3 b 1 | 9.6±0.5 b 12 | 3.3±0.2 1 | 3.2±0.6 12 |
| | 42 | 6.5±0.2 a 1 | 9.8±0.3 b 1 | 10.3±0.4 b 2 | 3.3±0.2 1 | 3.8±0.4 2 |
| | 46 | 6.7±0.4 a 1 | 8.4±0.6 a 1 | n.d. | 1.6±0.8 1 | n.d. |
| FW, mg | 24 | 89±4 a 1 | 114±5 b 1 | 110±3 b 1 | 25±8 1 | 21±6 1 |
| | 37 | 91±3 a 1 | 96±3 a 2 | 98±3 a 2 | 6±3 12 | 8±6 1 |
| | 42 | 88±4 a 1 | 107±3 b 1 | 98±3 a 2 | 18±5 1 | 9±6 1 |
| | 46 | 95±5 a 1 | 81±6 a 3 | n.d. | -14±11 2 | n.d. |

"Stem" — stem with leaf sheaths.
All designations are the same as in Tables S1.

**Table S6.** Water content (%) in plant organs before the experiment (7 day) and after growth under control or elevated temperatures at contrast conditions of air relative humidity (9 day).

| | Temp. | *Zea mays* | | | *Hordeum vulgare* | | |
|---|---|---|---|---|---|---|---|
| | | Plant age | | | Plant age | | |
| | °C | 7-day | 9-day | | 7-day | 9-day | |
| | | | HH | LH | | HH | LH |
| $2^{nd}$ leaf | 24 | 91.4±0.1 a1 | 90.6±0.2 b1 | 91.0±0.2 ab1 | 89.3±0.8 a1 | 87.5±0.4 a1 | 88.7±0.3 a1 |
| | 37 | 91.4±0.1 a1 | 89.4±0.1 b2 | 89.8±0.2 b2 | 90.2±0.5 a1 | 87.3±0.1 b1 | 86.9±0.6 b2 |
| | 42 | 91.1±0.4 a1 | 88.8±0.3 b2 | 88.9±0.4 b2 | 90.5±0.4 a1 | 87.1±0.2 b1 | 86.0±0.7 b2 |
| | 46 | 91.4±0.2 a1 | 87.2±0.5 b3 | 82.5±0.3 c3 | 89.7±0.1 a1 | 71.9±1.5 b2 | n.d. |
| $1^{st}$ leaf | 24 | 91.3±0.2 a1 | 91.5±0.2 a1 | 91.5±0.3 a1 | 90.5±0.3 a1 | 90.1±0.2 a1 | 90.3±0.2 a1 |
| | 37 | 91.3±0.2 a1 | 91.1±0.2 a12 | 91.2±0.2 a1 | 90.8±0.3 a1 | 89.8±0.1 b1 | 89.7±0.4 ab1 |
| | 42 | 91.6±0.6 a1 | 90.2±0.4 a23 | 90.3±0.3 a1 | 90.6±0.2 a1 | 86.1±0.9 b2 | 85.6±0.8 b2 |
| | 46 | 91.2±0.2 a1 | 89.8±0.4 b3 | 86.8±0.2 c2 | 90.6±0.1 a1 | 71.4±2.6 c3 | 83.8±0.6 b2 |
| Stem | 24 | 94.2±0.1 a1 | 93.5±0.2 b1 | 93.9±0.2 b1 | 92.3±0.4 a1 | 92.0±0.1 a1 | 92.1±0.3 a1 |
| | 37 | 94.3±0.1 a1 | 92.5±0.2 b2 | 92.7±0.1 b2 | 92.9±0.2 a1 | 90.0±0.1 b3 | 90.3±0.4 b2 |
| | 42 | 94.2±0.1 a1 | 91.5±0.3 b3 | 91.6±0.5 b2 | 92.6±0.2 a1 | 90.8±0.1 b2 | 89.4±0.1 c2 |
| | 46 | 94.1±0.2 a1 | 92.3±0.1 b2 | 92.3±0.4 b2 | 92.9±0.1 a1 | 89.5±0.5 b12 | n.d. |
| Shoot | 24 | 92.8±0.1 a1 | 92.2±0.1 b1 | 92.6±0.1 a1 | 91.1±0.4 a1 | 90.2±0.2 a1 | 90.5±0.2 a1 |
| | 37 | 92.8±0.1 a1 | 91.2±0.2 b2 | 91.5±0.1 b2 | 91.5±0.3 a1 | 89.4±0.1 b2 | 89.4±0.4 b2 |
| | 42 | 92.7±0.2 a1 | 90.4±0.2 b3 | 90.4±0.2 b3 | 91.5±0.2 a1 | 88.0±0.5 b2 | 87.1±0.4 b3 |
| | 46 | 92.7±0.2 a1 | 90.3±0.2 b3 | 88.3±0.2 c4 | 91.4±0.1 a1 | 80.2±1.4 b3 | 54.5±4.9 c4 |
| Roots | 24 | 94.2±0.2 a1 | 93.5±0.1 b1 | 93.8±0.5ab12 | 90.8±0.4 a1 | 91.4±0.1 a1 | 91.5±0.1 a1 |
| | 37 | 94.1±0.1 a1 | 93.7±0.2 a1 | 94.1±0.1 a1 | 91.5±0.2 a1 | 90.1±0.4 b2 | 90.3±0.3 b2 |
| | 42 | 93.9±0.2 a1 | 93.0±0.1 b2 | 93.0±0.2 b2 | 90.6±0.4 a1 | 89.2±0.2 b2 | 89.6±0.1 b3 |
| | 46 | 93.8±0.1 a1 | 92.6±0.2 b2 | 92.7±0.1 b2 | 91.3±0.2 a1 | 90.2±0.5 a2 | 89.0±0.1 b4 |

Leaves – leaf blades, stem - stem with leaf sheaths, shoot – whole shoot.
All designations are the same as in Tables S1.

**Table S7.** Uptake of water-based mineral media from pots during 48 h treatment in the thermostat chambers, ml per single plant (see Methods).

| Temp. °C | *Hordeum vulgare* | | *Zea mays* | |
|---|---|---|---|---|
| | HH | LH | HH | LH |
| 24 | 2.4±0.2 a 1 | 3.8±0.8 a 1 | 2.3±0.4 a 1 | 3.5±0.6 a 1 |
| 37 | 8.5±0.5 b 1 | 10.9±0.4 b 2 | 9.7±1.2 b 1 | 10.4±1.0 b 1 |
| 42 | 8.0±0.7 b 1 * | 11.9±0.7 b 2 | 10.0±0.5 b 1* | 12.6±0.6 bc 2 |
| 46 | 8.0±0.4 b 1 | 10.4±0.8 b 2 * | 10.8±1.2 b 1 | 14.4±0.8 c 2 * |

a-c – differences between variants of temperature regime are significant at $p \leq 0.05$.
1-2 – difference between plants of one species at a same temperature and the different conditions of relative air humidity (HH & LH) is significant at $p \leq 0.05$.
* - difference between two species in same conditions (°C, RH %) is significant at $p \leq 0.05$.
HH – higher (relative) humidity of air, LH - lower (relative) humidity of air.
Data are presented as means ± standard error (SE).

**Table S8.** Contents of the photosynthetic pigments (µg/g FW) in the 1st and 2nd leaves before the experiment (7 day) and after growth at different temperatures and RH conditions (9 day).

| | Te | *Zea mays*, plant age | | | *Hordeum vulgare*, plant age | | |
|---|---|---|---|---|---|---|---|
| | mp. | 7-day | 9-day | | 7-day | 9-day | |
| | °C | | HH | LH | | HH | LH |
| 1st leaves | | | | | | | |
| Chl *a* | 24 | 1831±80 a1 | 1853±88 a1 | 1923±53 a1 | 1208±14 a1 | 1269±81 a1 | 1211±43 a12 |
| | 37 | 1794±74 a1 | 1979±69 a1 | 1856±32 a1 | 1215±35 a1 | 1326±72 a1 | 1294±56 a1 |
| | 42 | 1920±56 a1 | 1892±39 a1 | 1772±38 a1 | 1228±3 a1 | 1018±56 c2 | 1091±38 c2 |
| | 46 | 1817±80 a1 | 1959±123 a1 | 1551±50 c2 | 1243±33 a1 | 883±64 c2 | n.d. |
| Chl *b* | 24 | 481±13 a1 | 463±22 a1 | 470±16 a1 | 338±11 a1 | 334±25 a12 | 319±13 a2 |
| | 37 | 435±15 a1 | 478±8 b1 | 449±13 ab1 | 322±10 a1 | 381±20 b1 | 372±11 b1 |
| | 42 | 469±47 a1 | 446±11 a1 | 430±12 a1 | 345±3 a1 | 292±11 c23 | 315±13 ac2 |
| | 46 | 455±37 ac1 | 481±40 a1 | 376±8 c2 | 324±25 a1 | 259±17 a3 | n.d. |
| Chl *a/b* | 24 | 3,8±0,1 c2 | 4,0±0,1 ac1 | 4,1±0,1 a1 | 3,6±0,1 a12 | 3,8±0,1 a1 | 3,8±0,0 a1 |
| | 37 | 4,1±0,0 a1 | 4,1±0,1 a1 | 4,2±0,1 a1 | 3,8±0,0 a1 | 3,5±0,1 c2 | 3,5±0,1 c2 |
| | 42 | 4,2±0,4 a12 | 4,3±0,1 a1 | 4,1±0,0 a1 | 3,6±0,0 a2 | 3,5±0,1 a2 | 3,5±0,1 a2 |
| | 46 | 4,1±0,2 a12 | 4,1±0,1 a1 | 4,1±0,1 a1 | 3,9±0,2 a12 | 3,4±0,1 a2 | n.d. |
| Chl *a+b* | 24 | 2312±93 a1 | 2317±110 a1 | 2392±67 a1 | 1547±17 a1 | 1603±106 a12 | 1530±56 a12 |
| | 37 | 2229±89 a1 | 2456±76 a1 | 2304±41 a1 | 1536±45 a1 | 1707±92 a1 | 1666±66 a1 |
| | 42 | 2389±101 a1 | 2338±44 a1 | 2202±50 a1 | 1573±2 a1 | 1311±66 c23 | 1406±50 c2 |
| | 46 | 2272±114 a1 | 2440±162 a1 | 1927±52 c2 | 1567±58 a1 | 1142±81 c3 | n.d. |
| Car | 24 | 276±14 a1 | 280±10 a1 | 289±6 a1 | 205±5 a1 | 212±20 a1 | 193±6 a1 |
| | 37 | 269±17 a1 | 309±14 a1 | 285±10 a1 | 218±17 a1 | 229±13 a1 | 219±13 a1 |
| | 42 | 284±6 a1 | 292±11 a1 | 282±12 a1 | 199±7 a1 | 201±14 a1 | 210±5 a1 |
| | 46 | 276±12 a1 | 311±18 a1 | 246±5 c2 | 215±8 a1 | 187±14 a1 | n.d. |
| Chl/ Car | 24 | 8,4±0,1 a1 | 8,3±0,1 a1 | 8,3±0,2 a1 | 7,6±0,1 a1 | 7,6±0,2 ab1 | 7,9±0,1 b1 |
| | 37 | 8,3±0,2 a1 | 8,0±0,2 a1 | 8,1±0,3 a1 | 7,1±0,4 a1 | 7,5±0,1 a1 | 7,6±0,3 a1 |
| | 42 | 8,4±0,4 a1 | 8,0±0,2 a1 | 7,8±0,2 a1 | 7,9±0,3 a1 | 6,6±0,2 c2 | 6,7±0,1 c2 |
| | 46 | 8,2±0,3 a1 | 7,8±0,2 a1 | 7,9±0,2 a1 | 7,3±0,5 a1 | 6,1±0,1 c2 | n.d. |
| 2nd leaves | | | | | | | |
| Chl *a* | 24 | 1685±182 a12 | 1998±67 ab2 | 2196±50 b1 | 1035±63 a1 | 1603±64 b1 | 1609±31 b1 |
| | 37 | 1589±88 a2 | 2339±39 b1 | 2138±103 b1 | 905±77 a1 | 1574±61 b1 | 1603±80 b1 |
| | 42 | 1935±56 a1 | 2087±145 a12 | 1796±68 a2 | 1019±59 a1 | 1272±82 b2 | 1326±46 b2 |
| | 46 | 1704±65 a2 | 1728±163 a2 | 1297±59 c3 | 990±26 a1 | 977±99 a2 | n.d. |
| Chl *b* | 24 | 447±44 a1 | 489±11 a2 | 529±16 a1 | 304±16 a1 | 459±32 b12 | 460±11 b1 |
| | 37 | 392±20 a1 | 564±6 b1 | 525±43 b12 | 251±21 a1 | 480±19 b1 | 497±30 b1 |
| | 42 | 482±46 a1 | 475±25 a2 | 421±13 a2 | 282±41 a1 | 378±20 a2 | 381±11 a2 |
| | 46 | 423±38 a1 | 424±46 a2 | 322±23 a3 | 261±20 a1 | 271±25 a3 | n.d. |
| Chl *a/b* | 24 | 3,8±0,1 c2 | 4,1±0,1 a1 | 4,2±0,1 a1 | 3,4±0,1 a2 | 3,5±0,1 a1 | 3,5±0,1 a1 |
| | 37 | 4,1±0,0 a1 | 4,2±0,1 a1 | 4,1±0,2 a1 | 3,6±0,0 a1 | 3,3±0,1 c1 | 3,2±0,1 c2 |
| | 42 | 4,1±0,4 a12 | 4,4±0,1 a1 | 4,3±0,1 a1 | 3,8±0,5 a12 | 3,4±0,1 a1 | 3,5±0,1 a12 |
| | 46 | 4,1±0,3 a12 | 4,1±0,2 a1 | 4,1±0,2 a1 | 3,9±0,2 a12 | 3,6±0,2 a1 | n.d. |
| Chl *a+b* | 24 | 2132±226 a12 | 2487±78 ab2 | 2725±66 b1 | 1339±78 a1 | 2061±96 b1 | 2069±38 b1 |
| | 37 | 1981±107 a2 | 2903±36 b1 | 2663±145 b1 | 1156±98 a1 | 2054±79 b1 | 2101±110 b1 |
| | 42 | 2417±100 a1 | 2562±169 a12 | 2218±80 a2 | 1301±98 a1 | 1649±101 b2 | 1707±54 b2 |
| | 46 | 2127±100 a12 | 2152±205 a2 | 1620±80 c3 | 1251±45 a1 | 1248±122 a3 | n.d. |
| Car | 24 | 263±25 a1 | 309±15 ab2 | 333±7 b1 | 191±4 a1 | 275±14 b1 | 273±11 b1 |
| | 37 | 262±25 a1 | 362±14 b1 | 325±8 b1 | 183±20 a1 | 286±12 b1 | 290±17 b1 |
| | 42 | 297±12 a1 | 326±27 a12 | 284±21 a1 | 201±7 a1 | 249±16 b12 | 255±10 b1 |
| | 46 | 267±9 a1 | 278±27 ac2 | 220±12 c2 | 195±6 a1 | 211±21 a2 | n.d. |
| Chl/ Car | 24 | 8,1±0,2 a1 | 8,1±0,2 a1 | 8,2±0,1 a1 | 7,0±0,3 a1 | 7,5±0,1 a1 | 7,6±0,2 a1 |
| | 37 | 7,7±0,4 a1 | 8,0±0,2 a1 | 8,2±0,4 a12 | 6,4±0,2 a1 | 7,2±0,1 b2 | 7,3±0,2 b1 |
| | 42 | 8,2±0,6 a1 | 7,9±0,2 a1 | 7,9±0,4 a12 | 6,5±0,6 a1 | 6,6±0,1 a3 | 6,7±0,1 a2 |
| | 46 | 8,0±0,4 a1 | 7,8±0,3 a1 | 7,4±0,1 a2 | 6,5±0,4 a1 | 5,9±0,1 a4 | n.d. |

Chl – chlorophyll. Car – carotenoids. All other designations are the same as in Tables S1.

**Table S9.** Parameters of $P_{700}$ light absorption (Pm) and Chl fluorescence (Fm, Fv, Fo) in dark-adapted state after 48 h under diverse temperatures and air RH conditions (see Methods).

| | Species & RH | Temperature | | | |
|---|---|---|---|---|---|
| | | 24°C | 37°C | 42°C | 46°C |
| Fo | Zm HH | 0,440±0,020 ab 1 | 0,315±0,019 a 2 | 0,297±0,015 bc 2 | 0,274±0,023 a 2 |
| | Zm LH | 0,460±0,014 a 1 | 0,346±0,018 a 2 | 0,283±0,018 c 3 | 0,239±0,019 a 3 |
| | Hv HH | 0,407±0,018 b 1 | 0,325±0,014 a 23 | 0,351±0,022 ab 12 | 0,280±0,024 a 3 |
| | Hv LH | 0,414±0,019 ab 1 | 0,308±0,013 a 2 | 0,359±0,021 a 1 | 0,225±0,016 a 3 |
| Fm | Zm HH | 2,129±0,085 a 1 | 1,421±0,080 a 2 | 1,314±0,074 a 2 | 0,848±0,077 a 3 |
| | Zm LH | 2,218±0,060 a 1 | 1,555±0,078 a 2 | 1,235±0,082 a 3 | 0,605±0,053 b 4 |
| | Hv HH | 2,298±0,101 a 1 | 1,524±0,062 a 2 | 1,352±0,083 a 2 | 0,883±0,070 a 3 |
| | Hv LH | 2,352±0,114 a 1 | 1,468±0,067 a 2 | 1,357±0,080 a 2 | 0,629±0,039 b 3 |
| Fv | Zm HH | 1,690±0,066 b 1 | 1,106±0,062 a 2 | 1,017±0,059 a 2 | 0,574±0,056 a 3 |
| | Zm LH | 1,758±0,049 ab 1 | 1,209±0,060 a 2 | 0,952±0,064 a 3 | 0,365±0,040 b 4 |
| | Hv HH | 1,891±0,083 ab 1 | 1,199±0,048 a 2 | 1,001±0,062 a 3 | 0,602±0,047 a 4 |
| | Hv LH | 1,939±0,095 a 1 | 1,160±0,054 a 2 | 0,997±0,061 a 2 | 0,405±0,030 b 3 |
| Fv/Fm | Zm HH | 0,794±0,002 b 1 | 0,778±0,004 bc 2 | 0,771±0,003 a 3 | 0,669±0,012 ab 4 |
| | Zm LH | 0,792±0,003 b 1 | 0,778±0,002 c 2 | 0,769±0,004 a 2 | 0,590±0,023 c 3 |
| | Hv HH | 0,823±0,002 a 1 | 0,787±0,003 ab 2 | 0,740±0,005 b 3 | 0,683±0,010 a 4 |
| | Hv LH | 0,824±0,001 a 1 | 0,789±0,002 a 2 | 0,734±0,007 b 3 | 0,636±0,019 bc 4 |
| Pm | Zm HH | 1,063±0,051 a 1 | 1,041±0,053 a 1 | 0,794±0,055 a 2 | 0,483±0,075 a 3 |
| | Zm LH | 0,953±0,061 a 1 | 0,960±0,084 a 1 | 0,736±0,041 a 2 | 0,225±0,040 b 3 |
| | Hv HH | 0,957±0,048 a 1 | 0,728±0,041 b 2 | 0,776±0,057 a 2 | 0,430±0,050 a 3 |
| | Hv LH | 0,985±0,050 a 1 | 0,739±0,049 b 2 | 0,708±0,055 a 2 | 0,249±0,032 b 3 |
| Pm/Fv | Zm HH | 0,645±0,039 a 1 | 0,874±0,054 a 2 | 0,785±0,082 a 12 | 0,788±0,062 a 12 |
| | Zm LH | 0,558±0,035 ab 1 | 0,727±0,073 ab 2 | 0,719±0,050 a 2 | 0,769±0,105 a 12 |
| | Hv HH | 0,483±0,024 b 1 | 0,571±0,025 b 2 | 0,707±0,035 a 3 | 0,677±0,071 a 23 |
| | Hv LH | 0,488±0,029 b 1 | 0,619±0,040 b 2 | 0,671±0,040 a 2 | 0,632±0,057 a 2 |

Values Pm, Fm, Fv, Fo are in V units. RH – relative humidity of air.
a-c – differences between plants of a same temperature regime – Hv HH, Hv LH, Zm HH, Zm LH - are significant at $p \leq 0.05$.
Zm – *Zea mays*, Hv – *Hordeum vulgare*,
All other designations are the same as in Tables S1.

**Table S9** v2. Alternative representation of statistical analysis for selected parameters. Means ± SE are the same.

| | Species & RH | Temperature | | | |
|---|---|---|---|---|---|
| | | 24°C | 37°C | 42°C | 46°C |
| Fm | Zm HH | 2,129±0,085 m | 1,421±0,080 nop | 1,314±0,074 op | 0,848±0,077 q |
| | Zm LH | 2,218±0,060 m | 1,555±0,078 n | 1,235±0,082 p | 0,605±0,053 r |
| | Hv HH | 2,298±0,101 m | 1,524±0,062 n | 1,352±0,083 nop | 0,883±0,070 q |
| | Hv LH | 2,352±0,114 m | 1,468±0,067 no | 1,357±0,080 nop | 0,629±0,039 r |
| Fv | Zm HH | 1,690±0,066 m | 1,106±0,062 opq | 1,017±0,059 pq | 0,574±0,056 r |
| | Zm LH | 1,758±0,049 mn | 1,209±0,060 o | 0,952±0,064 q | 0,365±0,040 s |
| | Hv HH | 1,891±0,083 mn | 1,199±0,048 o | 1,001±0,062 pq | 0,602±0,047 r |
| | Hv LH | 1,939±0,095 n | 1,160±0,054 op | 0,997±0,061 pq | 0,405±0,030 s |
| Fv/Fm | Zm HH | 0,794±0,002 n | 0,778±0,004 op | 0,771±0,003 p | 0,669±0,012 su |
| | Zm LH | 0,792±0,003 n | 0,778±0,002 p | 0,769±0,004 p | 0,590±0,023 v |
| | Hv HH | 0,823±0,002 m | 0,787±0,003 no | 0,740±0,005 q | 0,683±0,010 s |
| | Hv LH | 0,824±0,001 m | 0,789±0,002 n | 0,734±0,007 q | 0,636±0,019 uv |
| Pm | Zm HH | 1,063±0,051 m | 1,041±0,053 m | 0,794±0,055 n | 0,483±0,075 o |
| | Zm LH | 0,953±0,061 m | 0,960±0,084 m | 0,736±0,041 n | 0,225±0,040 p |
| | Hv HH | 0,957±0,048 m | 0,728±0,041 n | 0,776±0,057 n | 0,430±0,050 o |
| | Hv LH | 0,985±0,050 m | 0,739±0,049 n | 0,708±0,055 n | 0,249±0,032 p |

m-v – differences between all variants (temperatre, air humidity, and species) are significant at $p \leq 0.05$.

For Fv/Fm, letters “r” and “t” were omitted because of bad visibility in Arial in Fig 5C.
