## Supplementary Figures for "Lower air humidity reduced both the plant growth and activities of photosystems I and II under prolonged heat stress"

Figure S1. The dynamics of Fv/Fm (alternative representation of data given in Fig. 5C).

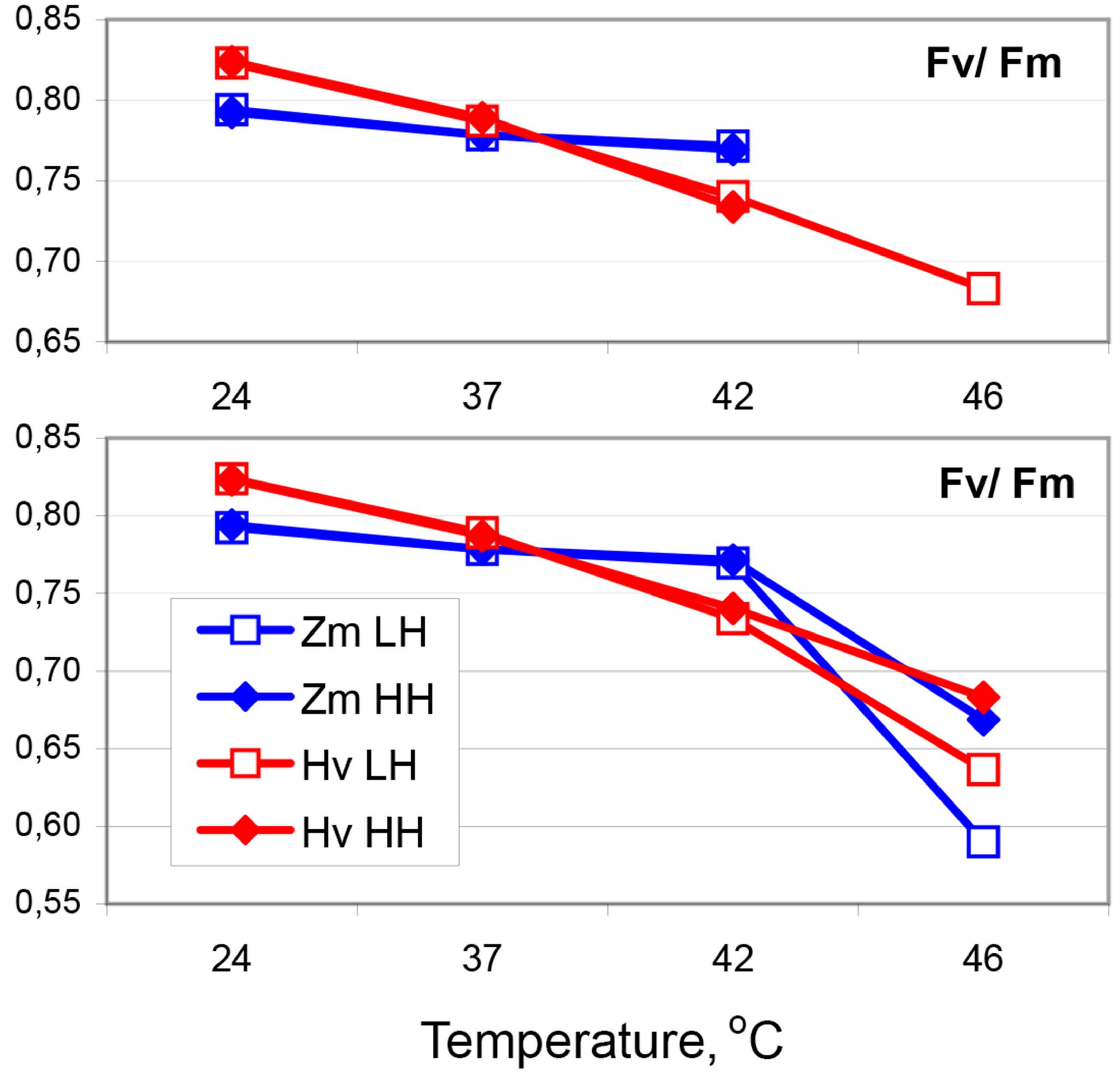


The upper panel shows incomplete data set to focus on the linear tendencies.
The lower panel shows complete data set.
SE bars are not shown (see Fig 5C).
Hv - Hordeum vulgare, Zm - Zea mays,
HH & LH - Higher & Lower relative humidity of air.

Figure S2. Dynamics of $\Phi_{PSII}$ in induction curves (IC).

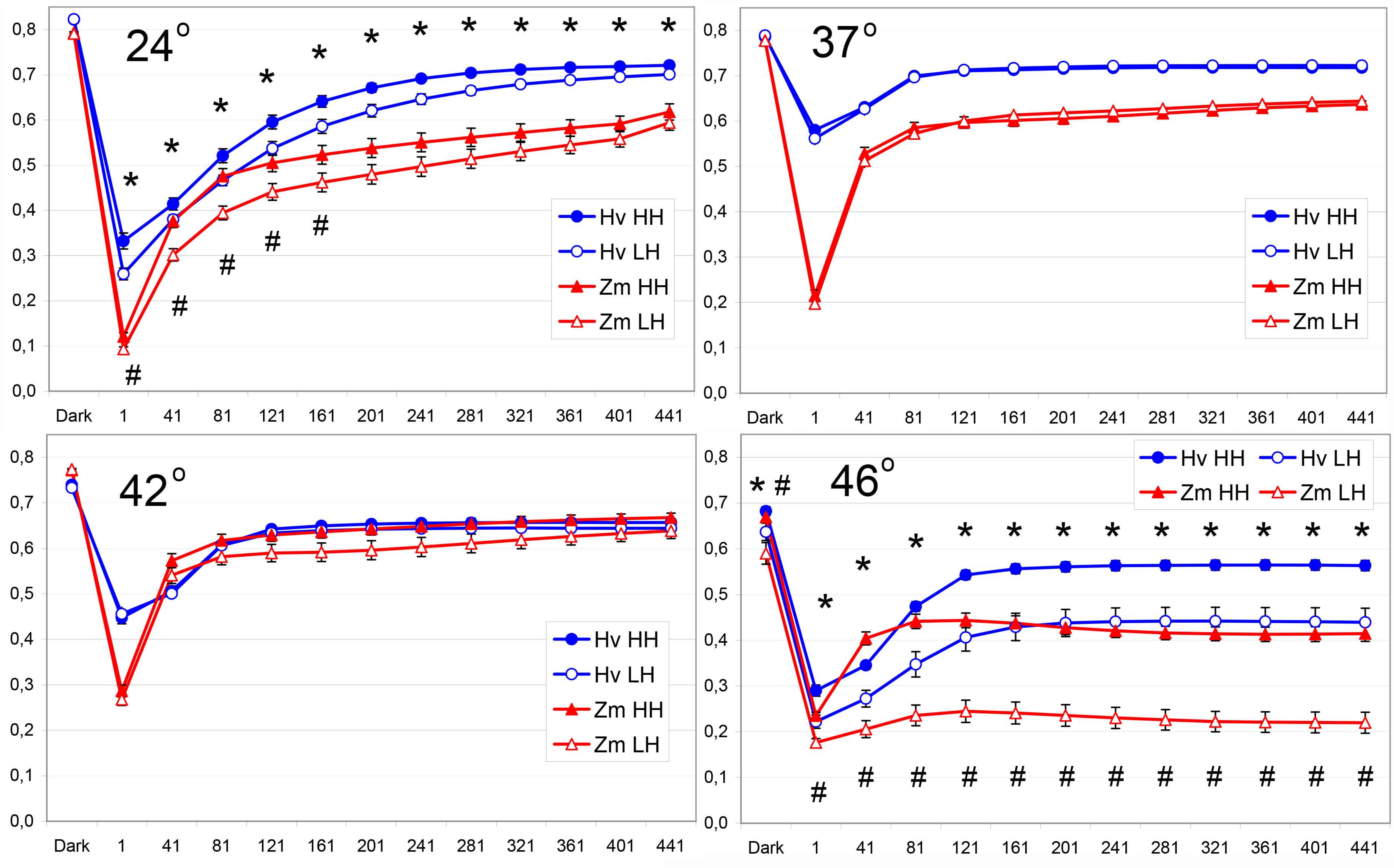


Time (AL), s

Hv - Hordeum vulgare, Zm - Zea mays, HH & LH - Higher & Lower relative humidity of air.

AL - actinic light, 70 µmol photons/ $m^2$ s. Temperature (°), Celsius degree.

*, # - significant difference between plants grown at different relative humidity of air (of the same species and at the same temperature), p<0.05; * - for Hv, # – for Zm.

Figure S3. Dynamics of the coefficients estimating photochemical component ($\Delta F$ = Fm' - Fs) of modulated Chl fluorescence: qP, $\Phi_{PSII}$, and X(II). Induction curves (IC). Revisualization of the data from Fig. 6 and Supplementary Fig. S2

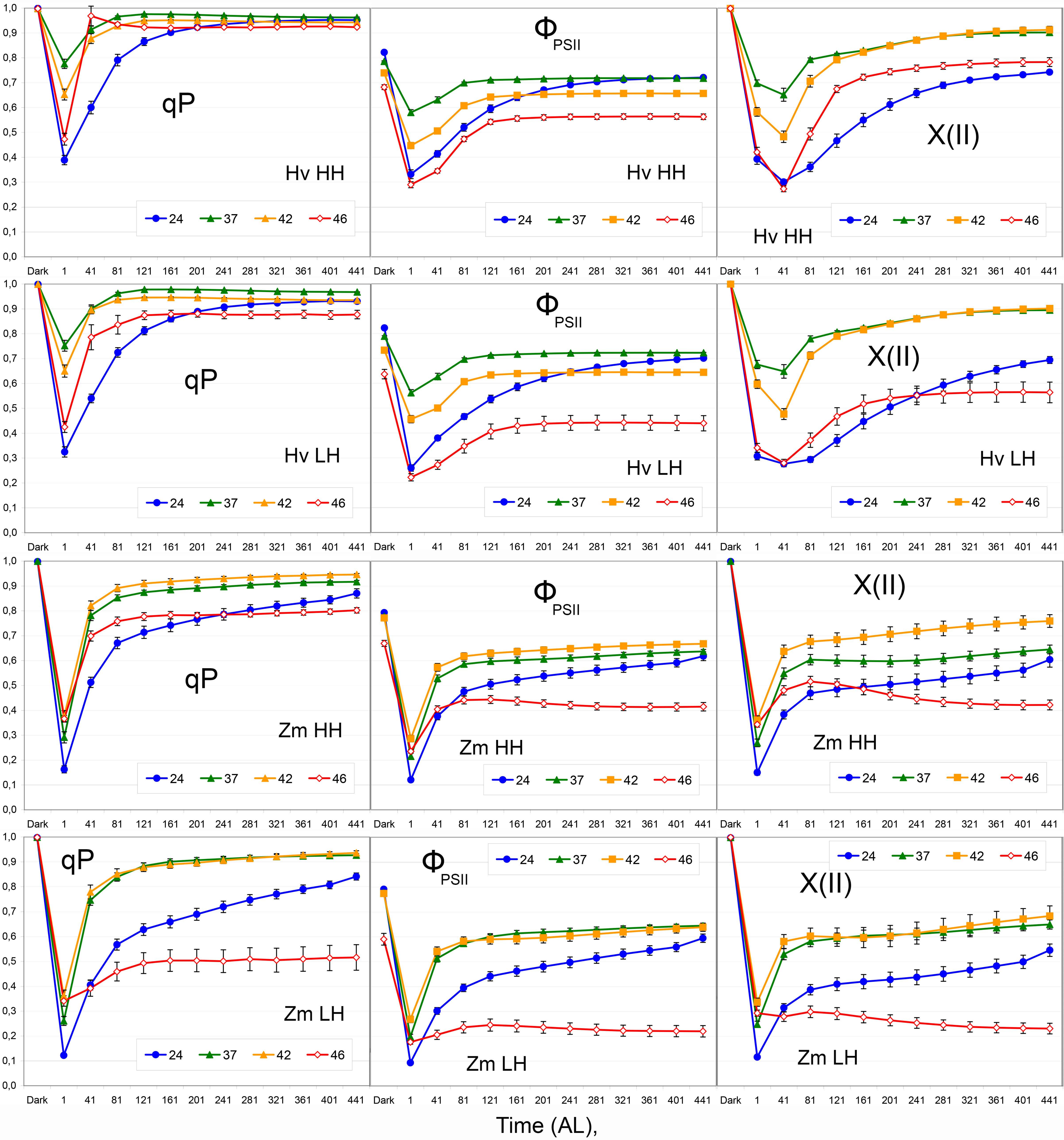


Numerals indicate temperature in Celsius degree. Coefficients qP = $\Delta F$/Fv', $\Phi_{PSII}$ = $\Delta F$/Fm', X(II) = $\Delta F$/Fv (see text). All other designations are the same as in Fig. S2.

### Figure S4. Dynamics of $\Phi_{PSII}$ in rapid light curves (RLC).

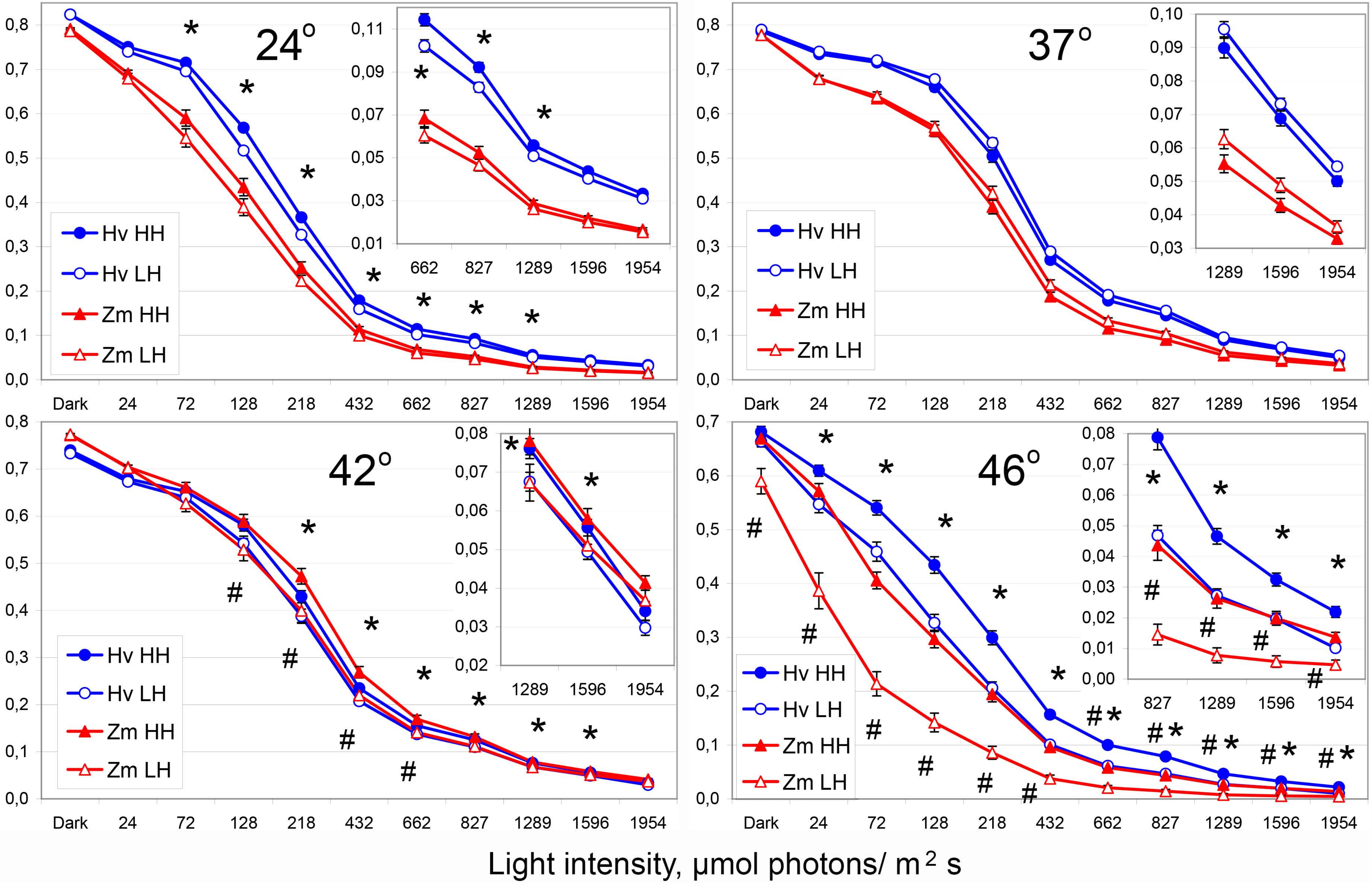


Insets demonstrate the corresponding small values with higher resolution.
All designations are the same as in Fig. S2.

Figure S5. Dynamics of the coefficients estimating photochemical component ($\Delta F = Fm' - Fs$) of modulated Chl fluorescence: qP, $\Phi_{PSII}$, and X(II). Rapid light curves (RLC). Revisualization of the data from Fig. 7 and Supplementary Fig. S4.

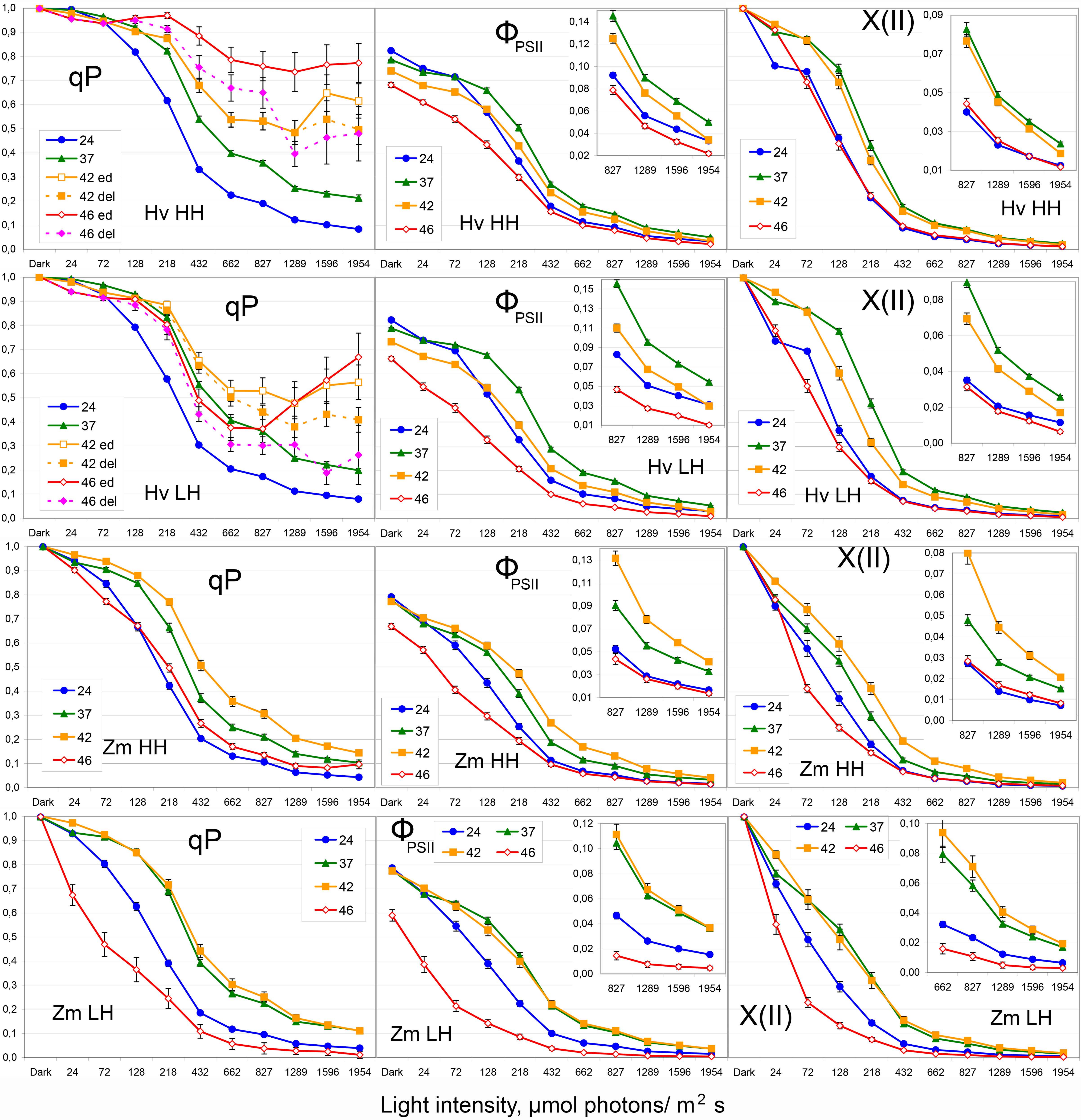


Numerals indicate temperature in Celsius degree. Coefficients $qP = \Delta F/Fv'$, $\Phi_{PSII} = \Delta F/Fm'$, $X(II) = \Delta F/Fv$ (see text). All other designations are the same as in Fig. S2.

Insets demonstrate the corresponding small values with higher resolution.

### Figure S6. Dynamics of Y(I) the quantum yield of PSI measured with IC (left) and RLC (right) methods. Revisualization of the data from Fig. 8.

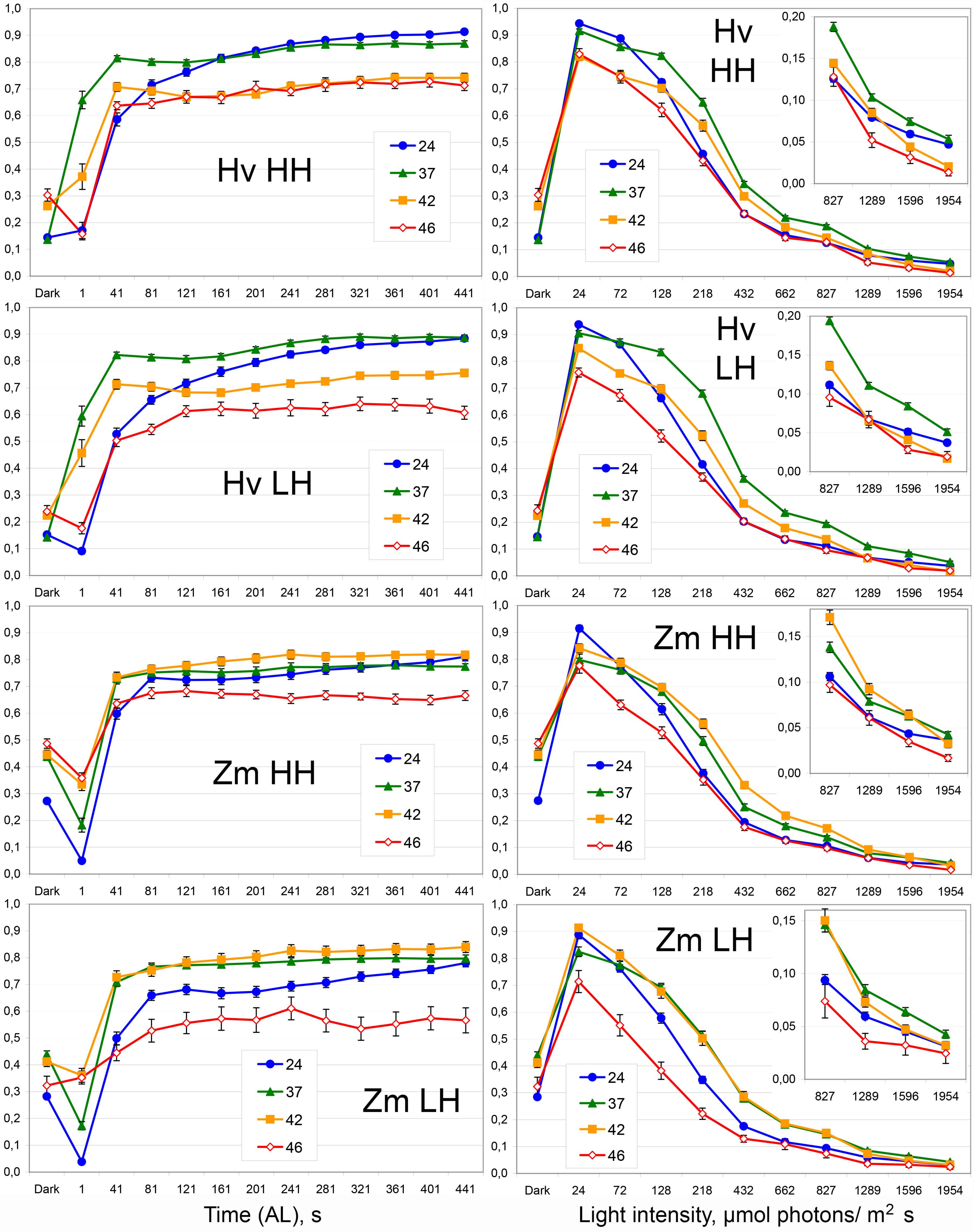


The induction curves (IC) are presented at left side; the rapid light curves (RLC) are presented at right side. Insets demonstrate the corresponding small values with higher resolution.
All designations are the same as in Fig. S2 and Fig. S4.

### Figure S7. The balance between photochemical processes in PSI and PSII calculated from the data of IC (left) and RLC (right). Revisualization of the data from Fig. 9.

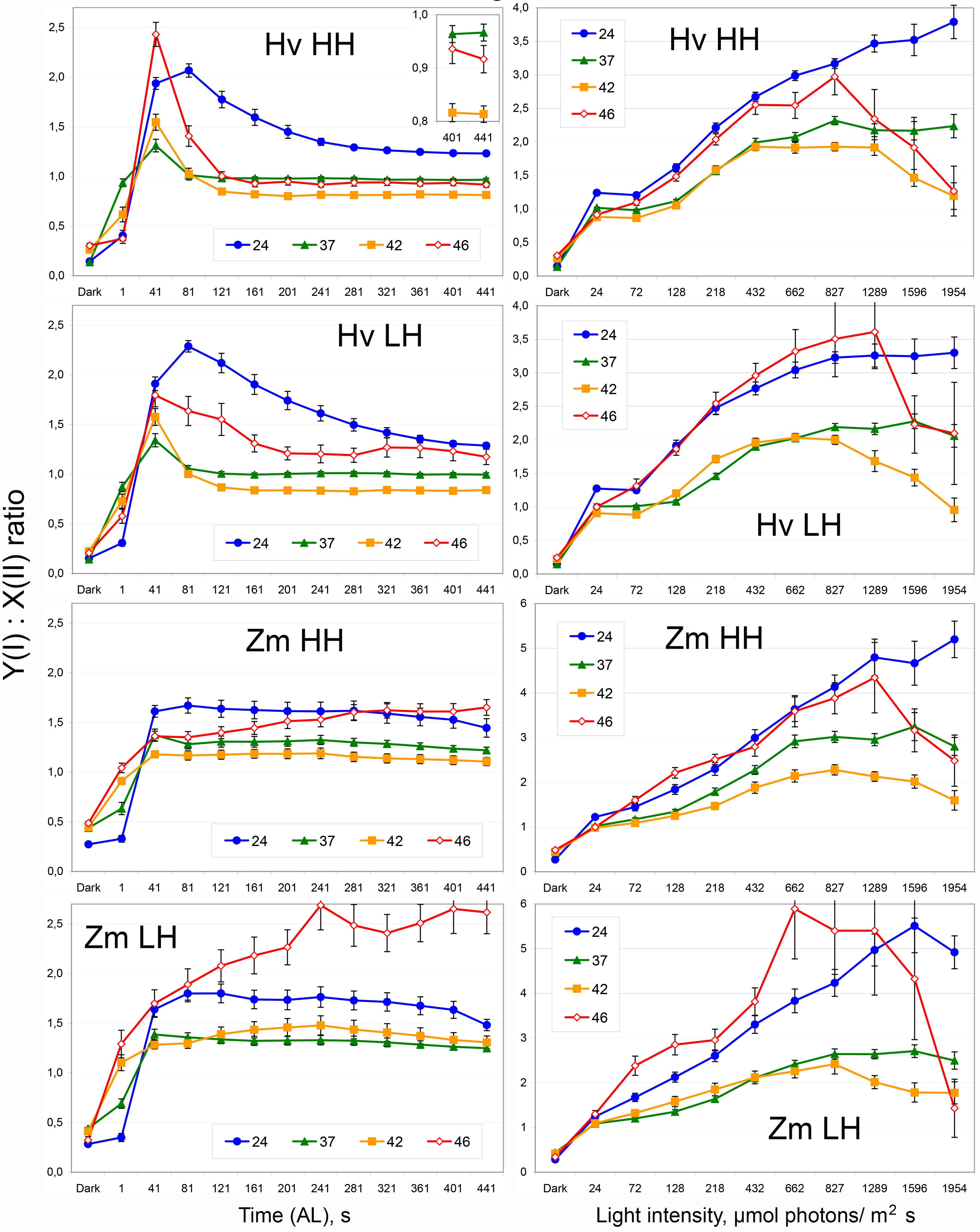


The induction curves (IC) are presented at left side; the rapid light curves (RLC) are presented at right side. Inset demonstrates the corresponding data with higher resolution.
All designations are the same as in Fig. S2 and Fig. S4.
